# Drug mechanism enrichment analysis improves prioritization of therapeutics for repurposing

**DOI:** 10.1101/2022.03.15.484520

**Authors:** Belinda B. Garana, James H. Joly, Alireza Delfarah, Hyunjun Hong, Nicholas A. Graham

## Abstract

**BACKGROUND:** There is a pressing need for improved methods to identify effective therapeutics for disease. Many computational approaches have been developed to repurpose existing drugs to meet this need. However, these tools often output long lists of candidate drugs that are difficult to interpret, and individual drug candidates may suffer from unknown off-target effects. We reasoned that an approach which aggregates information from multiple drugs that share a common mechanism of action (MOA) would increase on-target signal compared to evaluating drugs on an individual basis. In this study, we present Drug Mechanism Enrichment Analysis (DMEA), an adaptation of Gene Set Enrichment Analysis (GSEA), which groups drugs with shared MOAs to improve the prioritization of drug repurposing candidates.

**RESULTS:** First, we tested DMEA on simulated data and showed that it can sensitively and robustly identify an enriched drug MOA. Next, we used DMEA on three types of rank-ordered drug lists: (1) perturbagen signatures based on gene expression data, (2) drug sensitivity scores based on high-throughput cancer cell line screening, and (3) molecular classification scores of intrinsic and acquired drug resistance. In each case, DMEA detected the expected MOA as well as other relevant MOAs. Furthermore, the rankings of MOAs generated by DMEA were better than the original single-drug rankings in all tested data sets. Finally, in a drug discovery experiment, we identified potential senescence-inducing and senolytic drug MOAs for primary human mammary epithelial cells and then experimentally validated the senolytic effects of EGFR inhibitors.

**CONCLUSIONS:** DMEA is a fast and versatile bioinformatic tool that can improve the prioritization of candidates for drug repurposing. By grouping drugs with a shared MOA, DMEA increases on-target signal and reduces off-target effects compared to analysis of individual drugs. DMEA is publicly available as both a web application and an R package at https://belindabgarana.github.io/DMEA.

## BACKGROUND

Identifying effective therapeutics for disease remains a pressing challenge. Recent efforts in large-scale ‘omic profiling [1–4], pharmacological and genetic loss-of-function screening [5–7], and drug perturbation profiling [8] have generated a wealth of molecular data characterizing large numbers of cell lines and their responses to perturbations. Many computational approaches have been developed to leverage these molecular data for drug sensitivity predictions and/or drug repurposing [9–25], and these efforts have successfully identified drugs for a wide variety of diseases, including HIV [18], osteoporosis [26], diabetes [27], and cancer [14, 28, 29]. Despite these successes, many patients remain ineligible for targeted therapies, including over 80% of cancer patients [30]. Furthermore, only about half of eligible cancer patients are responsive to targeted therapy, emphasizing the need for improved drug discovery and repurposing methods.

One common drawback of many drug repurposing tools is that they output a long list of candidate drugs with limited information about how the top candidates are related. This complicates efforts to prioritize drugs on the list for validation, as researchers must consider many drugs targeting different molecular pathways with the caveat that some targeted therapies may not actually inhibit their intended target [31]. Therefore, given a list of candidate drugs, we reasoned that grouping drugs with similar mechanisms of action (MOAs) into a “set” and then statistically evaluating the enrichment of the drug set in the list would increase on-target signal and reduce off-target effects compared to evaluating drugs on an individual basis. Our approach, called Drug Mechanism Enrichment Analysis (DMEA), is an adaptation of the popular Gene Set Enrichment Analysis (GSEA) algorithm [32] in which drugs, rather than genes, are grouped into sets based on annotated MOAs. Each drug set is then statistically evaluated against a background of all other drug sets. If multiple drugs which share a common MOA are all highly ranked candidates, then this indicates that the identified MOA is more likely to be a true on-target sensitivity.

Notable alternatives to our approach for analyzing enriched MOA in drug lists include the Connectivity Map (CMap) [8] and Drug Set Enrichment Analysis (DSEA) [25]. However, these tools have several key limitations (Fig. 1) including that they: 1) can only query CMap’s L1000 reduced representation transcriptomic database; 2) have limited statistical rigor (e.g., lack of p-values in CMap, lack of multiple hypothesis correction in DSEA); 3) accept only one type of unranked input list (i.e., gene names for CMap, drug names for DSEA); and 4) do not generate plots of the overall and MOA-specific results. In addition, we note that DSEA queries gene sets (e.g., gene ontology terms like “cellular protein localization”) rather than drug MOAs (e.g., HDAC inhibitors). To address these shortcomings, we sought to make DMEA compatible with any dataset or drug repurposing algorithm, maintain the statistical rigor of GSEA, and generate plots of the overall results (i.e., volcano plot of all MOA normalized enrichment scores) and MOA-specific results (i.e., mountain plots). Furthermore, to address a lack of web-accessible tools to identify selectively toxic drugs based on an input gene signature, we included a feature to pair DMEA with a simple molecular classification method (i.e., weighted gene voting [33]). This rapid procedure can be used with any number of genes, requires less than a minute to execute, and can be accessed online with no programming required.

**Figure 1.**
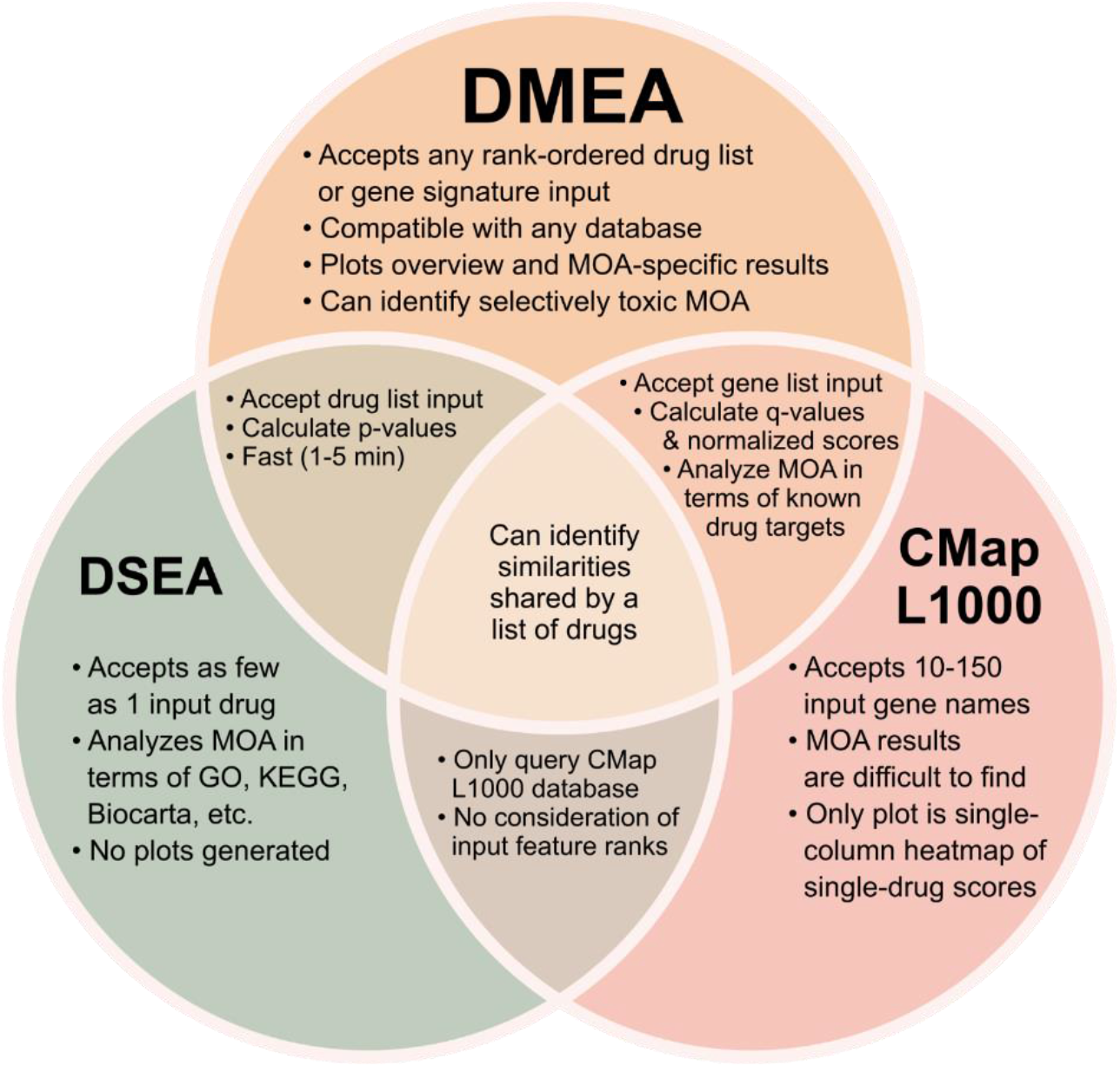
DMEA is more flexible and statistically rigorous than other approaches to evaluate drug MOA. The Venn diagram compares our method, DMEA, with those of Drug-Set Enrichment Analysis (DSEA) [25] and Connectivity Map (CMap) L1000 query of gene expression signatures [8].

In summary, DMEA can help researchers better prioritize potential drug treatments by aggregating results across many drugs which share MOAs to identify global trends. By quantifying the coordinated enrichment of drugs annotated with the same MOA and normalizing scores across a large background of drug MOAs, DMEA can improve on-target prioritization of candidates for drug repurposing. DMEA is publicly available as a web application or an R package at https://belindabgarana.github.io/DMEA.

## METHODS

### Drug Mechanism Enrichment Analysis (DMEA)

DMEA tests whether drugs known to share a MOA are enriched in a rank-ordered drug list. DMEA can be applied to any rank-ordered list of drugs with annotations for at least two MOAs. For a drug MOA to be evaluated, at least six drugs (or the minimum number of drugs per MOA set by the user) must be annotated with that MOA and each drug must be ranked by a nonzero numeric value. DMEA uses the same algorithm as GSEA [32] but applies it to sets of drugs, rather than genes, to identify drug MOAs which are overrepresented at either end of the input rank-ordered drug list (further detail below). If a drug MOA is positively enriched, then drugs annotated with that MOA are overrepresented at the top of the list. Conversely, if a drug MOA is negatively enriched, then drugs which share that MOA annotation are overrepresented at the bottom of the list.

Specifically, for each MOA, DMEA calculates an enrichment score (ES) as the maximum deviation from zero of a running-sum, weighted Kolmogorov-Smirnov-like statistic. The p-value is estimated using an empirical permutation test wherein drugs are randomly assigned MOA labels in 1,000 independent permutations to calculate a distribution of null enrichment scores (ES_null); the p-value is then calculated as the percentage of same-signed ES_null equal to or greater than the ES divided by the percentage of same-signed ES_null. The normalized enrichment score (NES) is then calculated by dividing the ES by the mean of the same-signed portion of the ES_null distribution. Finally, the q-value or false discovery rate (FDR) is calculated as the percentage of same-signed NES in the null distribution (i.e., NES_null) with NES equal or greater to the observed NES divided by the percentage of same-signed NES equal or greater. We use a significance threshold of p < 0.05 and FDR < 0.25 by default per the recommendation for GSEA, but this FDR cutoff can be customized by the user. Given a rank-ordered drug list, DMEA generates 1) enrichment results for all tested drug MOAs; 2) a volcano plot summarizing the NES and −log_10_(p-value) for all tested drug MOAs; and 3) mountain plot(s) for individual drug MOA(s) which pass the given FDR cutoff (Fig. 2).

**Figure 2.**
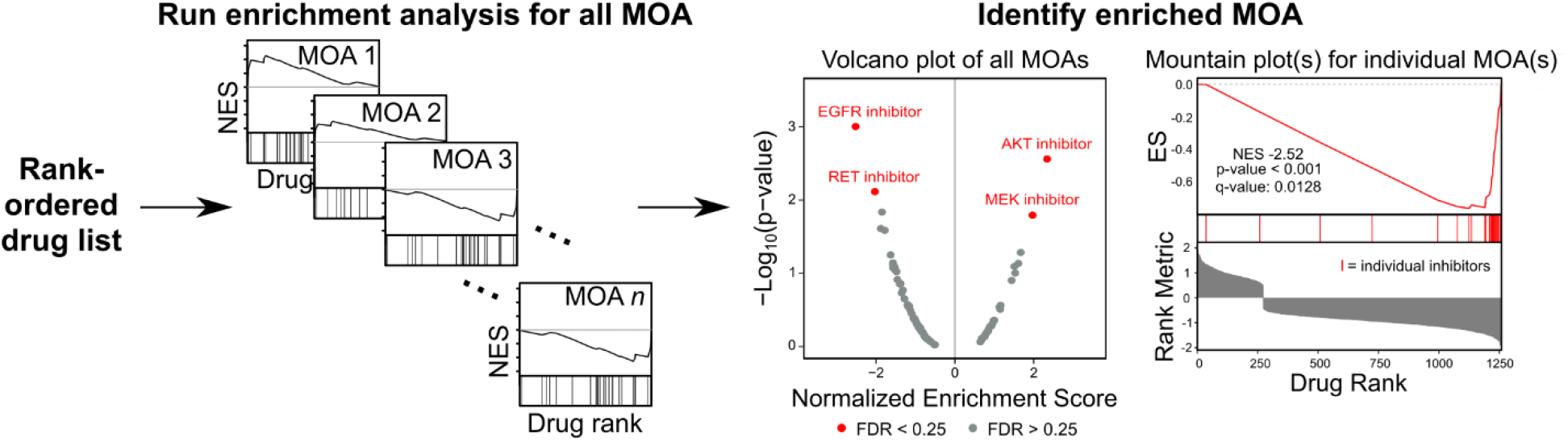
Overview of Drug Mechanism Enrichment Analysis. DMEA is an adaptation of GSEA which analyzes a rank-ordered drug list to identify drug MOAs that are overrepresented at either end of the input drug list. Given a rank-ordered drug list where drugs have been annotated with known MOAs, DMEA runs an enrichment analysis for each individual MOA. After calculating p-values and FDR q-values, DMEA outputs 1) enrichment results for all tested drug MOAs; 2) a volcano plot summarizing the NES and −log_10_(p-value) for all tested drug MOAs; and 3) mountain plot(s) for individual drug MOA(s) which pass the FDR cutoff.

### Simulation study of DMEA

To evaluate the sensitivity of DMEA, we first simulated a rank-ordered drug list by randomly assigning values from a normal distribution (μ = 0, σ = 0.5) for 1,351 drugs with MOA annotations in the PRISM drug screen. Next, a number of drugs, *X*, were randomly sampled as a synthetic drug set and their rank values were selected from a shifted normal distribution (μ = *Y*, σ = 0.5); the size of the synthetic drug set, *X*, was varied from 5 to 50 drugs, and the perturbation value *Y* was varied from −1 to +1. This rank-ordered drug list was then analyzed by DMEA to determine the enrichment of the synthetic drug set relative to the known drug MOA sets provided by the PRISM drug screen. For each pair of *X* and *Y* values, the simulation was repeated 50 times to assess reproducibility (i.e., the ability of DMEA to consistently detect a true difference between the synthetic drug set and the background drug sets determined by MOA annotations from PRISM).

### CMap L1000 Query

The Connectivity Map (CMap) web portal (https://clue.io) [8] allows users to query the L1000 gene expression database using 10 to 150 input up- and down-regulated gene IDs. The output is a normalized connectivity score which indicates the similarity between the query and differentially expressed gene sets induced by drug treatments. A positive score indicates similarity between the query and the perturbagen signature, whereas a negative score indicates dissimilarity. Specifically, we used the “query.gct” file from the zip file output by the CMap L1000 Query (found within their “/arfs/TAG” folder), including the MOA annotations provided in the file. Since this file includes results for all cell lines in the L1000 database as well as information for quality control, we removed any scores indicated to be low quality and averaged scores across cell lines for each drug with MOA annotations. Here, we used example data sets from the CMap web portal, including: 1) GSE32547, HUVEC cells treated with the HMGCR inhibitor pitavastatin (1 μM, 4h) or DMSO [34]; 2) GSE35230, A375 melanoma clones treated with the MEK inhibitor GSK212 (30 nM, 24 h) or DMSO [35]; 3) GSE14003, JEKO1 cells treated with the proteasome inhibitor bortezomib (10 h) or untreated [36]; 4) GSE28896, human CD34^+^ cells treated with the glucocorticoid agonist dexamethasone (24 h) or untreated [37]; and 5) GSE33643, A2058 cells treated with the PI3K/MTOR inhibitor BEZ235 (3 doses at 24 h) or DMSO [38]. We also used the up- and down-regulated biomarkers from a proteomic signature of senescence in primary human mammary epithelial cells (HMECs) [39]. To compare DMEA’s results to CMap’s MOA enrichment results, we used the “gsea_result.gct” file found within the “\gsea\TAG\arfs\NORM_CS” folder. We compared the results for chemical perturbagens combined across all cell lines, specifically “cell_iname = −666” with pert_type = “TRT_CP”, for either set_type = PCL” or “set_type = “MOA_CLASS”.

### CMap PRISM Query

The Connectivity Map (CMap) web portal (https://clue.io) [8], also allows users to query PRISM viability data [5] for 3 to 489 input cell line IDs classified as having “UP” or “DOWN” phenotypes. The query outputs a normalized connectivity score ranking the drugs based on their toxicity towards the “UP” versus “DOWN” cell lines. A positive score indicates toxicity towards “UP” cell lines, whereas a negative score indicates toxicity towards “DOWN” cell lines or lack of toxicity towards “UP” cell lines. In particular, we used the “ncs.gct” file from the zip file output by the CMap PRISM Query, including the MOA annotations provided in the file. Again, we only considered drugs with MOA annotations. Here, we used examples provided by the CMap web portal including: 1) cell lines with the EGFR activating mutation p.E746_A750del (i.e., “UP” cell lines: NCIH1650, PC14, and HCC827); 2) cell lines with high expression of *PDGFRA* (i.e., “UP” cell lines: 42MGBA, A204, A2780, G292CLONEA141B1, G402, GB1, HS618T, KNS42, LMSU, MG63, MON, NCIH1703, SBC5, SKNAS, SNU685, SW579, U118MG, U251MG, and YH13); and 3) cell lines sensitive to the HMGCR inhibitor lovastatin (i.e., “UP” cell lines: HUH28, SNU1079, MG63, LOXIMVI, MDAMB231, SF295, SNU1105, YKG1, ACHN, HCT15, SNU423, SNU886, CALU1, HCC4006, HCC44, HCC827, NCIH1915, NCIH661, NCIH838, PC14, RERFLCMS, SQ1, SW1573, KYSE150, A2780, COV434, JHOM1, MCAS, KP3, SW1990, MSTO211H, YD15, HS944T, MDAMB435S, MELJUSO, A204, HT1080, RH30, LMSU, FTC238, YD8, 5637, and AGS).

### Weighted gene voting (WGV)

To calculate a molecular classification score for cell lines based on external molecular signatures, we used weighted gene voting (WGV) [33]. The WGV score is the dot product between an external gene signature of interest and normalized RNAseq expression values for 327 adherent cancer cell lines from the Cancer Cell Line Encyclopedia (CCLE, version 19Q4) [3]. In this study, we analyzed four independent transcriptomic signatures, three derived from datasets for intrinsic resistance to EGFR inhibitors and one derived from a dataset for acquired resistance to a RAF inhibitor. For each transcriptomic data set, we used the R package limma [40] to perform an eBayes statistical analysis for differential expression comparing sensitive and resistant samples. Then, the top 500 genes based on |log_2_(fold-change)| with q-value < 0.05 were used for WGV (with the log_2_(fold-change) being the gene “weight” or rank value).

For gene signatures of EGFR inhibitor sensitivity, we used data sets GSE12790 [41], GSE31625 [42], and Coldren et al. [43]. In GSE12790, transcriptomic profiles were provided for breast cancer cell lines classified as either sensitive (EC50 < 1 µM: HDQ-P1, CAL85-1, and HCC1806) or resistant to erlotinib (EC50 > 10 µM: CAL-51, CAL-120, MDA-MB-231, BT-20, HCC1569, EFM-192A, HCC1954, MDA-MB-453, BT474, HCC1428, T47D, ZR-75-1, KPL-1, BT-483, MDA-MB-415, HCC1500, CAMA-1, and MCF7). For GSE31625, we used 17 transcriptomic profiles from 3 non-small cell lung cancer cell lines sensitive (H1650, H3255, and PC-9) and 12 profiles of 2 cell lines resistant to erlotinib (A549 and UKY-29). Finally, based on classifications from Coldren *et al*., we used CCLE RNAseq profiles of 5 non-small cell lung cancer cell lines sensitive (NCIH1650, HCC95, NCIH1975, NCIH1648, and NCIH2126) and 7 cell lines resistant to gefitinib (NCIH520, NCIH460, NCIH1299, HCC44, A549, NCIH1703, and HCC15). For a gene signature of RAF inhibitor sensitivity, we used dataset GSE66539 with paired biopsy samples of melanoma from 3 patients before vemurafenib treatment and after emergence of resistance to vemurafenib [44].

### DMEA using WGV molecular classification scores

For rapid identification of drug MOAs with toxicity towards cells represented by an input gene signature, DMEA can be used in combination with a molecular classification method such as WGV, correlations, and large public databases for gene expression and drug screens. To do this, we accessed the Cancer Cell Line Encyclopedia version 19Q4 for RNAseq data and calculated WGV scores for 327 adherent cancer cell lines using external molecular signatures. To avoid overfitting, we did not include WGV scores from any cell lines that had been used to generate the external molecular signature. Next, we calculated the Pearson correlation between the WGV scores and PRISM drug sensitivity scores (i.e., area under the curve (AUC) values for cell viability as a function of drug concentration) for each drug [5] using data from the most recent PRISM screen available (e.g., HTS002, MTS005, MTS006, and MTS010). Lastly, drugs were ranked by the Pearson correlation coefficient, and the rank-ordered drug list was analyzed by DMEA using the MOA annotations provided in the PRISM dataset.

### Simulation study of DMEA using WGV molecular classification scores

For 200 synthetic cell lines, we sampled drug sensitivity scores for 1,351 drugs with MOA annotations in the PRISM drug screen from a bimodal mixture of two normal distributions (μ_1_=0.83, σ_1_=0.08 and μ_2_=1.31, σ_2_=0.08) with the lower distribution containing 72% of all drugs. This distribution was chosen to reflect the distribution of the PRISM drug sensitivity data (i.e., AUC) [5]. Additionally, we simulated gene expression for each cell line by sampling from a normal distribution with a mean (μ) of 0 and standard deviation (σ) of 0.5. This distribution was chosen to reflect the distribution of the normalized CCLE RNAseq data [3].

To introduce a synthetic association between gene expression and drug sensitivity, we randomly sampled a synthetic gene set of 25 genes and a synthetic drug set of 10 drugs. Next, expression values for the synthetic gene set and sensitivity scores for the synthetic drug set were each sampled from a shifted distribution, where the magnitude of the shift for each synthetic cell line is determined by a perturbation value ranging from 0 (no perturbation) to 0.1. For example, for a perturbation value of 0.1, the mean gene expression for the 25 perturbed genes in cell line 1 was μ = −0.1, and the mean sequentially increased by 0.001 for cell lines 2-200; similarly, the mean drug sensitivity of cell line 1 to the 10 perturbed drugs was shifted by −0.1, and this shift value sequentially increased by 0.001 for cell lines 2-200. This created a gradient of perturbations in the 200 cell lines, meaning cell line 1 had the largest negative perturbation and cell line 200 had the largest positive perturbation. Then, we calculated WGV scores for each cell line by taking the dot product of the expression values of the synthetic gene set and the difference in average gene expression between the top and bottom 10 percent of cell lines (i.e., gene weights from cell lines 181-200 which had the highest mean expression versus cell lines 1-20 which had the lowest mean expression). Afterwards, we calculated the Pearson correlation between the WGV and drug sensitivity scores for each of the 1,351 drugs in the synthetic dataset. Finally, drugs were ranked by their Pearson correlation coefficient, and DMEA was performed to measure the enrichment of the synthetic drug set relative to the background drug sets which were determined by the MOA annotations in the PRISM drug screen. To assess reproducibility, this entire process was repeated 50 times.

### Cell culture

HMEC cells were purchased from Thermo Scientific and cultured in M87A medium (50% MM4 medium and 50% MCDB170 supplemented with 5 ng/ml EGF, 300 ng/ml hydrocortisone, 7.5 µg/ml insulin, 35 µg/ml BPE, 2.5 µg/ml transferrin, 5 µM isoproterenol, 50 µM ethanolamine, 50 µM o-phosphoethanolamine, 0.25 % FBS, 5 nM triiodothyronine, 0.5 nM estradiol, 0.5 ng/ml cholera toxin, 0.1 nM oxytocin, 1% anti-anti, no AlbuMax) in atmospheric oxygen. Glucose and glutamine-free DMEM was purchased from Corning (90-113-PB), Ham’s F12 was purchased from US Biological (N8542-12), and MCD170 medium was purchased from Caisson Labs (MBL04). Glucose and glutamine were added to the media at the appropriate concentration for each media type. At each passage, cells were lifted with TrypLE at 80-90% confluency and seeded at a density of 2.3 × 10^3^/cm^2^.

### Cell viability experiments

Proliferating HMECs (PD ∼12) were seeded at a concentration of 2.1 × 10^3^/cm^2^ or 9.5 × 10^3^/cm^2^ for DMSO and triapine treatment, respectively. The following day, cells were treated with DMSO (vehicle control) or 2 µM triapine for 3 days. The cells were counted and then treated with either DMSO (vehicle control), dacomitinib, AZD8931, or navitoclax at 100 nM or 500 nM for 3 days. Cell viability and live cell number were measured with trypan blue assay using a TC20 automated cell counter (Bio-Rad). Chemical inhibitors were from Sigma (triapine) or MedChemExpress (dacomitinib, AZD8931, navitoclax).

## RESULTS

### DMEA identifies an enriched drug MOA in simulated data

To evaluate the ability of DMEA to identify the enrichment of drug sets, we tested it on a normally distributed, synthetic ranked list of 1,351 drugs (see Methods). For a randomly sampled set of drugs ranging in size from 5 to 50 drugs, we shifted these drugs’ rankings by a perturbation value ranging from −1 to 1. Next, we ran DMEA using the full rank list of drugs to assess enrichment of the synthetic drug set. This process was repeated 50 times for each synthetic drug set size, after which the average normalized enrichment score (NES) and the percentage of replicates with significant enrichment of the synthetic drug set were visualized as heatmaps (Fig. 3).

**Figure 3.**
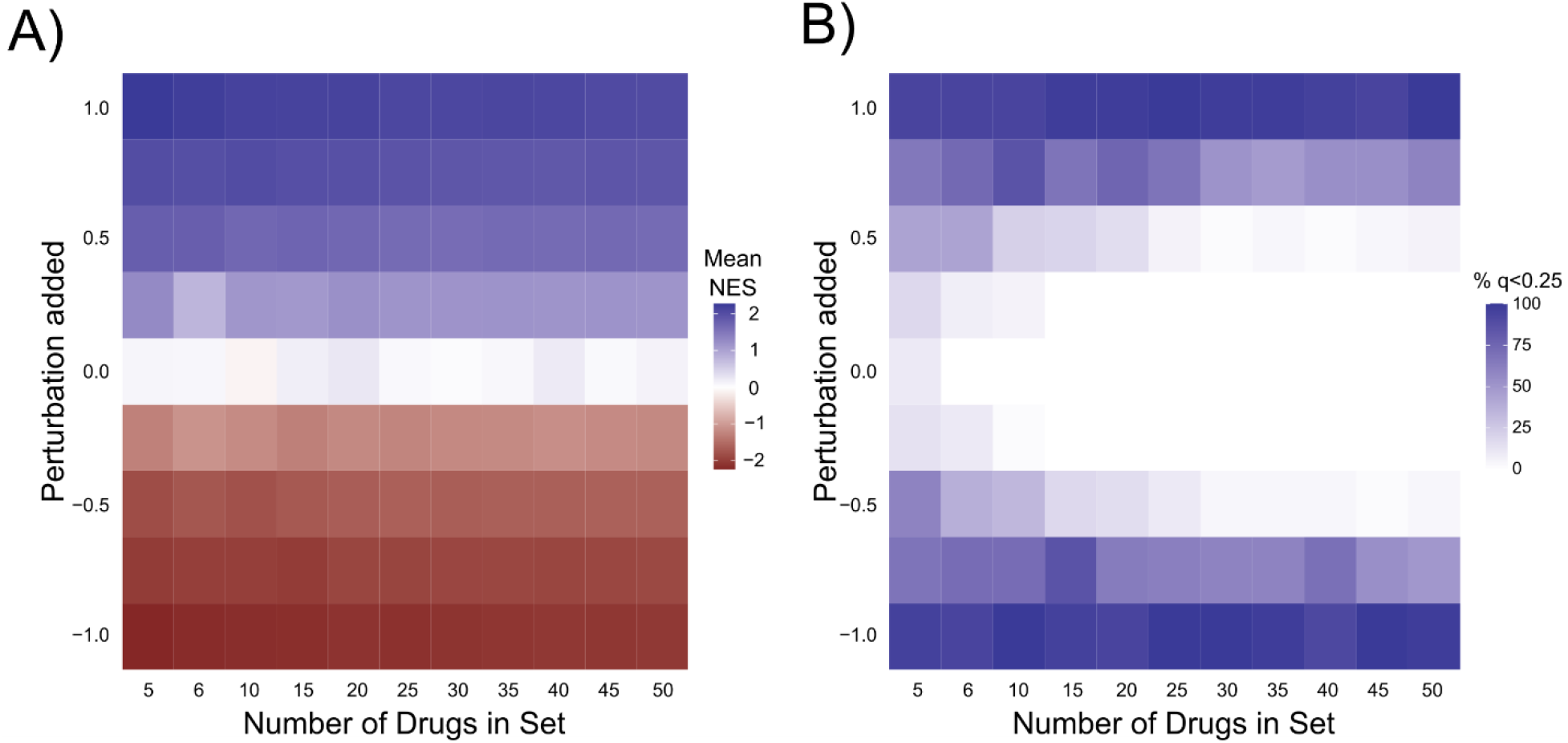
Sensitivity analysis of DMEA using synthetic data. Synthetic rank-ordered drug lists were generated with varying perturbations (y-axis) of different drug set sizes (x-axis), then analyzed by DMEA (see Methods). For each combination of drug set size and perturbation value, 50 replicates were performed. A) Heatmap showing the average DMEA NES for the perturbed drug set. B) Heatmap showing the percent of DMEA replicates with FDR q-value < 0.25 for the perturbed drug set.

As expected, we observed no false discovery except for very small drug sets (i.e., when evaluating a set of 5 drugs we observed 2% of replicates were falsely enriched). As the magnitude of the perturbation was increased or decreased, the average NES of the synthetic drug set increased or decreased, respectively. Likewise, the percentage of replicates passing the significance threshold of p < 0.05 and FDR < 0.25 increased as the magnitude of the perturbation increased. These results demonstrate that DMEA can successfully identify an enriched set of drugs in simulated data.

### DMEA identifies similar MOAs based on gene expression connectivity scores

Next, we sought to test whether DMEA could identify enriched drug MOAs in rank-ordered drug lists generated by the Connectivity Map (CMap), [8] a popular tool for drug discovery that contains >1 million gene expression signatures measured using L1000, a reduced representation transcriptomic profiling method. Specifically, we analyzed example data sets from the CMap L1000 Query tool to identify perturbagen signatures that are similar or dissimilar to an input gene set. First, we used a gene expression signature from HUVEC cells treated with pitavastatin, an inhibitor of 3-hydroxy-3-methylglutaryl-CoA reductase (HMGCR) [34], to rank 3,868 drugs based on the similarity of their effects on gene expression. Because pitavastatin itself was found in the list of 3,868 drugs, one might have expected it to be the top-ranked, most similar drug produced by this analysis, but in fact it ranked 24^th^ out of 3,868 drugs (Supp. Fig. 1A). In contrast, analysis of the rank-ordered list of drugs using DMEA identified the HMGCR MOA as the only significant similar MOA (Fig. 4A). This demonstrates that analysis of MOAs by DMEA can generate clearer and more statistically significant results that analysis of individual drugs in results from the CMap L1000 Query.

**Figure 4.**
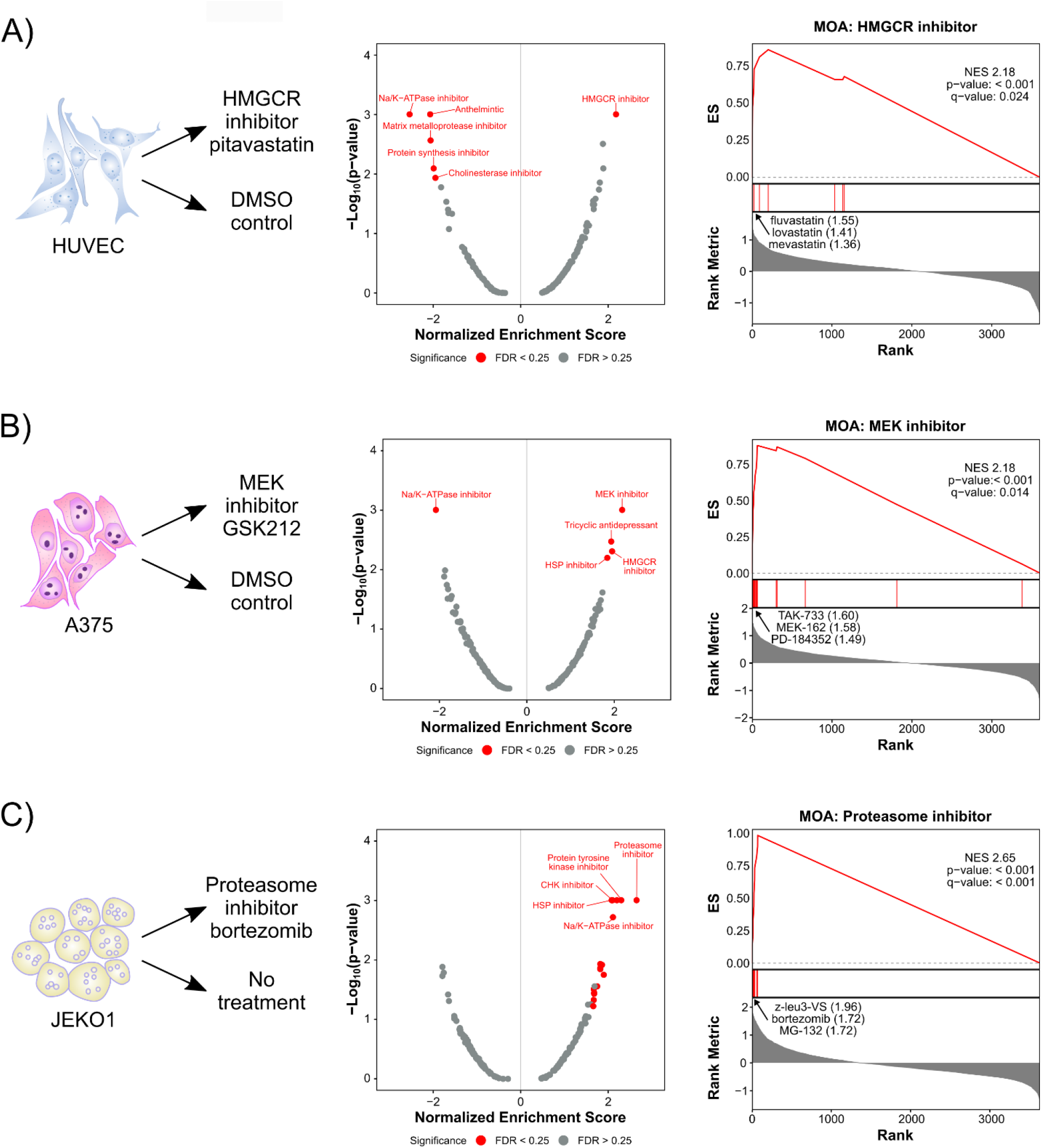
DMEA identifies similar MOAs based on gene expression connectivity scores. Rank-ordered drug lists were generated by querying the CMap L1000 gene expression perturbation signatures and then analyzed by DMEA. A) HUVEC cells treated with the HMGCR inhibitor pitavastatin [34], B) A375 melanoma clones treated with the MEK inhibitor GSK212 [35], and C) JEKO1 cells treated with the proteasome inhibitor bortezomib [36]. Volcano plots summarizing the NES and −log_10_(p-value) for all tested drug MOAs and mountain plots of the expected MOAs are shown. Red text indicates MOAs with p-value < 0.05 and FDR < 0.25. For each mountain plot, the inhibitors with the most positive connectivity scores are highlighted.

Next, we tested a gene expression signature from A375 melanoma cells treated with the MEK inhibitor GSK212 [35]. Again, DMEA correctly identified that MEK inhibitors were the most similar MOA in the rank-ordered list of drugs (Fig. 4B; Supp. Fig. 1A). In this case, comparison to the single drug rankings was not possible because the L1000 database does not contain the true drug treatment, GSK212 (Supp. Fig. 1A). Subsequently, we analyzed a gene expression signature from JEKO1 mantle cell lymphoma cells treated with the proteasome inhibitor bortezomib [36]. DMEA again accurately found that proteasome inhibitors were the most similar MOA (Fig. 4C), and DMEA’s MOA-level ranking (#1) was improved upon the single-drug ranking of the true drug treatment, bortezomib (#14/3,868) (Supp. Fig. 1A). Finally, we used DMEA to test data sets from human CD34^+^ cells treated with the glucocorticoid agonist dexamethasone [37] and A2058 melanoma cells treated with the PI3K/mTOR inhibitor BEZ235 [38]. In both cases, DMEA correctly identified the expected MOA as significantly enriched (glucocorticoid receptor agonist and PI3K/mTOR inhibitor MOAs, respectively) (Supp. Fig. 2) and DMEA’s MOA-level rankings were improved upon CMap’s individual drug rankings of the true drug treatments (Supp. Fig. 1A). Taken together, these results show that DMEA can correctly identify enriched MOAs in rank-ordered lists of drugs generated by the CMap L1000 Query, and that the MOA-level rankings of the true drug treatments are improved compared to the single-drug rankings.

Next, we compared DMEA’s MOA-level results to those of the CMap L1000 Query (found in an output sub-folder generated by CMap L1000 Query called “\gsea\TAG\arfs\NORM_CS”). Like our DMEA results, CMap’s MOA-level rankings revealed the expected MOA as the top-ranked most enriched MOA in all cases except for glucocorticoid receptor agonists and PI3K inhibitors which were not found in the L1000 output (Supp. Fig. 1A). We also compared our DMEA results to the CMap L1000’s perturbagen classes (PCLs), which are derived from MOA sets but exclude drugs which do not fit the overall trend of the MOA [8]. Again, CMap’s PCL rankings were similar to that of DMEA (Supp. Fig. 1A). Thus, DMEA and the CMap L1000 generate similar MOA-level rankings. However, in contrast to DMEA, the CMap L1000 MOA-based analysis has less statistical rigor (i.e., no p-values provided by CMap) and does not generate any plots of the overall and MOA-specific results (e.g., volcano or mountain plots).

### DMEA identifies selectively toxic MOAs based on cell viability connectivity scores

To evaluate if DMEA can identify enriched MOAs in a different type of rank-ordered drug list, we used the CMap PRISM Query tool to query data from the PRISM drug repurposing database [5]. Given an input list of cell line names, the CMap PRISM Query generates a list of ∼1,200 drugs ranked by normalized connectivity scores which represent the predicted sensitivity of the input cell lines to each drug. The higher the normalized connectivity score, the more toxic the drug is predicted to be for the input cell lines. Again, we analyzed example data sets from the CMap PRISM Query tool to test DMEA, including: 1) cell lines with the activating EGFR mutation p.E746_A750del (n=3), 2) cell lines with high expression of *PDGFRA* (n=19), and 3) cell lines with sensitivity to the HMGCR inhibitor lovastatin (n=43). As hypothesized, DMEA identified EGFR inhibitors (Fig. 5A), PDGFR inhibitors (Fig. 5B), and HMGCR inhibitors (Fig. 5C), respectively, as significantly positively enriched in these rank-ordered drug lists. DMEA also improved upon the rank of the true drug sensitivity for the HMGCR inhibitor lovastatin (#1 with DMEA’s MOA-level rankings versus #3 in the single-drug rankings, Supp. Fig. 1B). Altogether, these examples demonstrate that DMEA can identify enriched MOA in rank-ordered lists of drugs generated by CMap Query of the PRISM drug screen, and that the MOA-level ranking of the true drug sensitivity is higher than that of the single-drug ranking.

**Figure 5.**
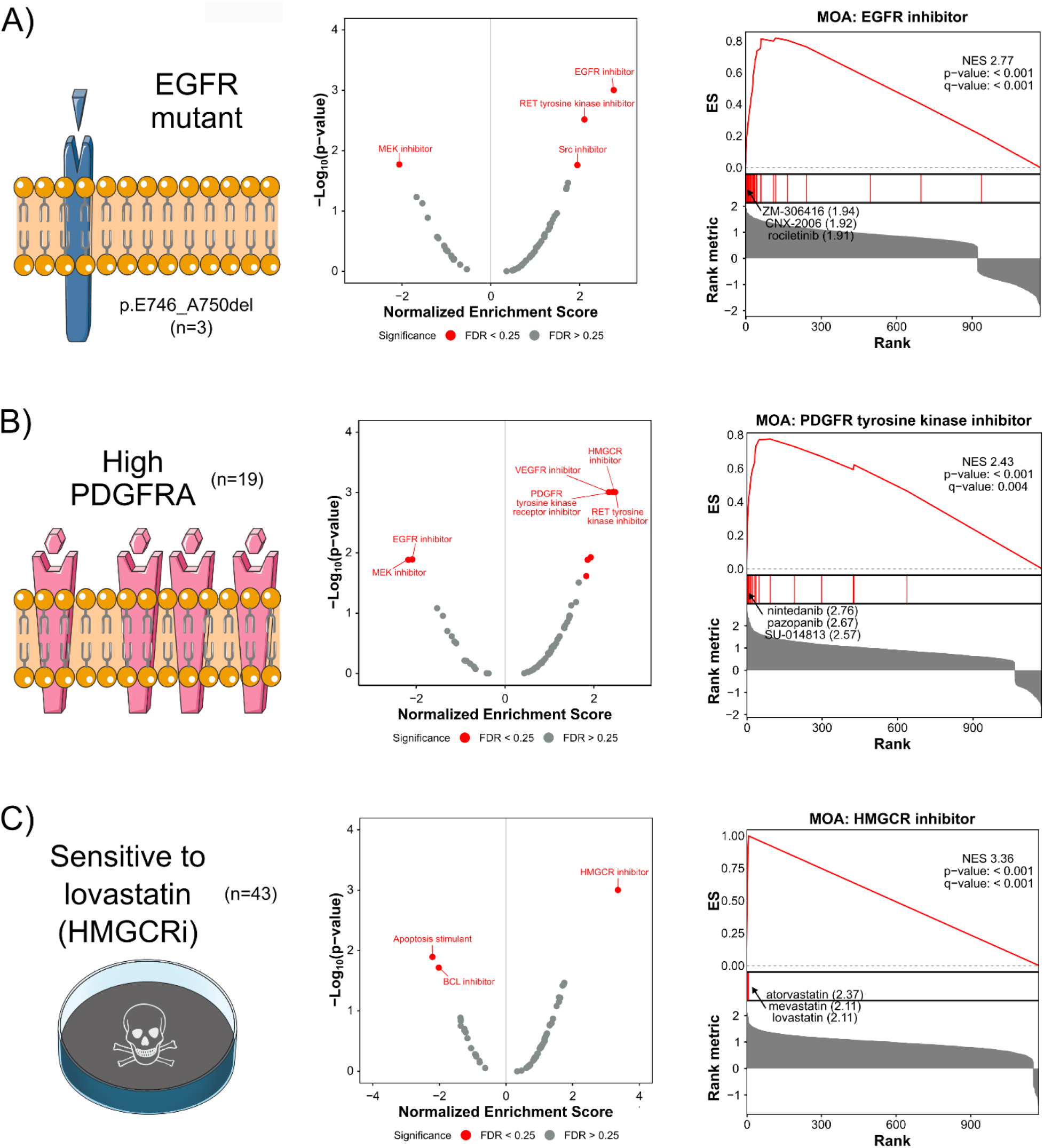
DMEA identifies selectively toxic MOAs based on cell viability connectivity scores. Rank-ordered drug lists were generated by querying the PRISM database with input cell line sets characterized by A) the activating EGFR mutation p.E746_A750del, B) high expression of *PDGFRA*, and C) sensitivity to the HMGCR inhibitor lovastatin. Volcano plots summarizing the NES and −log_10_(p-value) for all tested drug MOAs and mountain plots of the expected MOAs are shown. Red text indicates MOAs with p-value < 0.05 and FDR < 0.25. For each mountain plot, the inhibitors with the most positive connectivity scores are highlighted.

### DMEA identifies selectively toxic MOAs based on molecular signatures

To offer a web-accessible method to identify selectively toxic drug MOAs based on an input molecular signature (i.e., up- or down-regulated genes that characterize a disease or cell type), we paired DMEA with a simple molecular classification method, namely weighted gene voting (WGV) [33]. Specifically, we: 1) used WGV to classify adherent cancer cell lines in the CCLE database based on similarity to the input gene signature; 2) correlated WGV scores with drug sensitivity scores (i.e., AUC) for each of 1,351 drugs in the PRISM database; and 3) ranked drugs by the correlation coefficient of WGV scores and drug AUC values (Supp. Fig. 3; see Methods).

To test this approach, we first performed a simulation study. Specifically, we simulated gene expression and drug sensitivity scores for 200 cell lines by randomly sampling values from distributions that reflected the CCLE RNAseq and PRISM drug sensitivity data. Next, to create a synthetic association between gene expression and drug sensitivity, we perturbed a subset of the gene expression data and drug sensitivity scores. We then ran 50 replicates to determine if DMEA could consistently identify enrichment of the synthetic drug set in this simulated data. To visualize the results, we plotted the average normalized enrichment score (NES) and the percent of replicates which pass the significance threshold of FDR < 0.25 as heatmaps (Supp. Fig. 4). Importantly, when there was no perturbation in drug sensitivity (AUC), the tested drug set was never significantly enriched (0% of replicates) regardless of the size of the perturbation in RNA expression. This demonstrates that DMEA is not prone to false positive results using this WGV-based approach. In addition, increasing the perturbation in either RNA expression or drug sensitivity led to increased enrichment scores (i.e., average NES) and increased significance (i.e., higher percentage of significant replicates). These results illustrate that DMEA can successfully identify associations between gene expression and drug sensitivity with high reproducibility in simulated data.

Next, we tested whether DMEA could successfully identify drug MOAs with selective toxicity using published transcriptomic signatures of drug resistance. First, we tested three different signatures of intrinsic resistance to EGFR inhibitors: 1) non-small cell lung cancer (NSCLC) cell lines treated with erlotinib (GSE31625) [42]; 2) breast cancer cell lines treated with erlotinib (GSE12790) [41]; and 3) NSCLC cell lines treated with gefitinib (Coldren *et al*.) [43]. Notably, there was little overlap in the genes used for WGV (GSE12790 and Coldren *et al*. share zero genes, GSE12790 and GSE31625 share 15 genes, and GSE31625 and Coldren *et al*. share 19 genes; Fig. 6B). Despite the lack of overlap in input gene signatures, all three DMEA analyses correctly identified EGFR inhibitors as the top toxic drug MOA for the EGFR inhibitor-sensitive cancer cell lines (Fig. 6A-B, Supp. Fig. 5). Again, DMEA’s MOA-level rankings were improved compared to the single-drug rankings (#1 for EGFR inhibitors in all cases versus #16 for erlotinib based on GSE31625, #10 for erlotinib based on GSE12790, and #13 for gefitinib based on Coldren *et al*.; Supp. Fig. 1B). In addition, DMEA revealed consistent results across all three input gene signatures for drug MOAs identified as potentially toxic for EGFR inhibitor-resistant cancer cell lines, including HMGCR and MDM inhibitors (Fig. 6B, Supp. Fig. 5). These results support that DMEA can identify selectively toxic drug MOAs given a molecular signature of intrinsic drug resistance and that DMEA’s MOA-level rankings improve upon single-drug rankings of toxicity.

**Figure 6.**
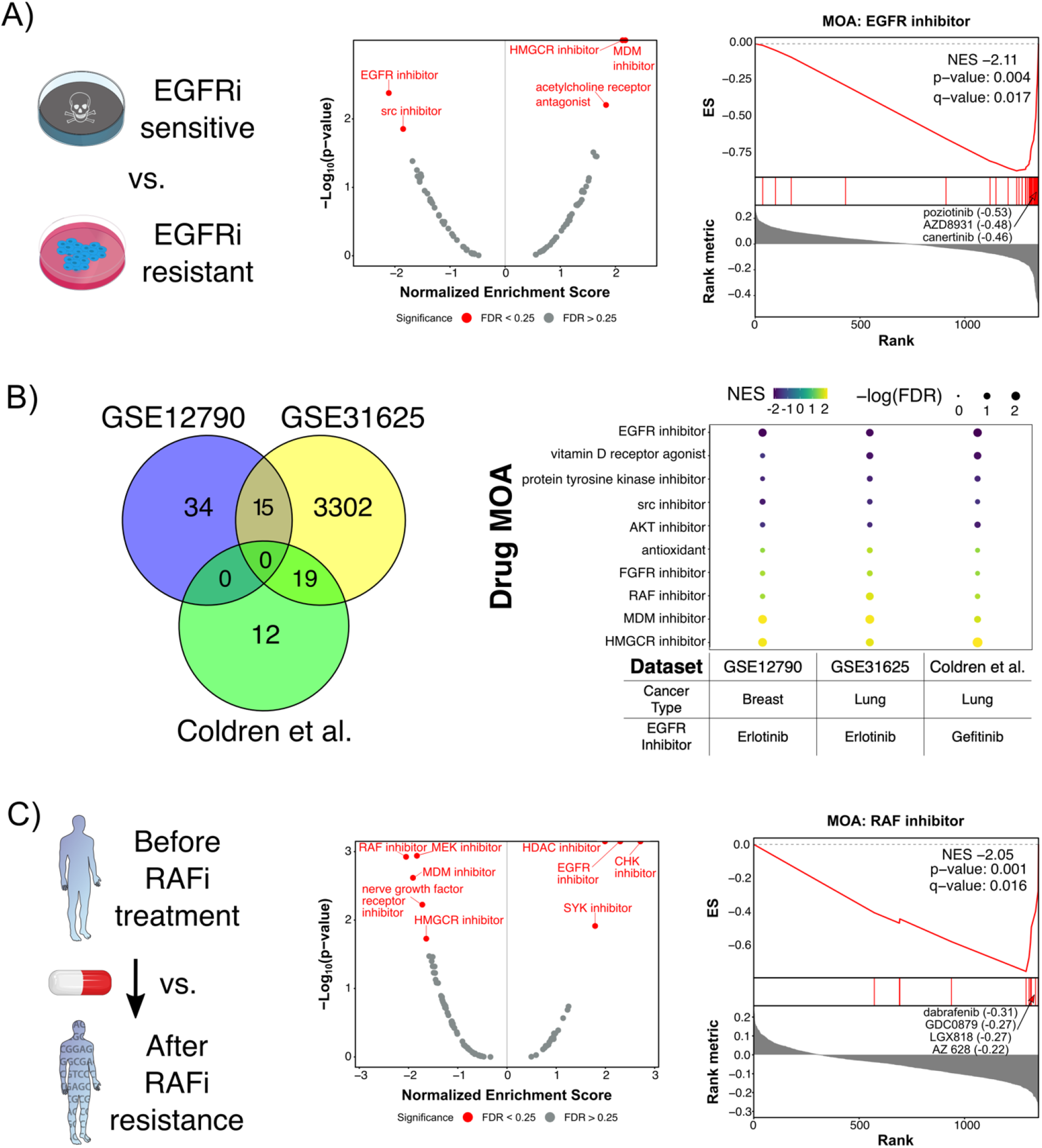
DMEA identifies selective toxic MOAs based on external gene expression signatures of intrinsic EGFR inhibitor resistance and acquired RAF inhibitor resistance, respectively. Using gene expression signatures of intrinsic resistance to EGFR inhibition and acquired resistance to RAF inhibition, we calculated WGV molecular classification scores for 327 adherent cancer cell lines in the CCLE database. For each signature, the WGV scores were correlated with drug sensitivity scores (i.e., AUC) for 1,351 drugs from the PRISM database. Drugs were then ranked by Pearson correlation coefficient, and DMEA was performed to identify selectively toxic MOAs. A) DMEA analysis of GSE12790 [41] transcriptomic signature of intrinsic resistance to EGFR inhibitor erlotinib, including a volcano plot of NES versus −log_10_(p-value) for MOA evaluated where red text indicates MOAs with p-value < 0.05 and FDR < 0.25 and a mountain plot showing that DMEA identified the EGFR inhibitor MOA as negatively enriched. The most negatively correlated EGFR inhibitors are labeled along with their correlation coefficients. B) Comparison of three transcriptomic signatures for intrinsic resistance to EGFR inhibition analyzed using DMEA, including a Venn diagram showing the number of shared genes among the signatures and a dot plot illustrating the consistency of MOA enrichment across DMEA’s analyses. C) DMEA analysis of GSE66539 [44] transcriptomic signature of acquired resistance to RAF inhibitor vemurafenib, including a volcano plot of NES versus −log_10_(p-value) for MOA evaluated where red text indicates MOAs with p-value < 0.05 and FDR < 0.25 and a mountain plot showing that DMEA identified the RAF inhibitor MOA as negatively enriched. The most negatively correlated RAF inhibitors are labeled along with their correlation coefficients.

Next, we tested whether DMEA could identify selectively toxic drug MOAs given a transcriptomic signature of acquired drug resistance. Specifically, we analyzed RNAseq data from patient biopsies of BRAF-mutant melanoma before treatment with the BRAF inhibitor vemurafenib and after the development of acquired resistance [44]. Again, DMEA correctly identified the RAF inhibitor MOA as the top toxic drug MOA for the samples collected prior to BRAF inhibitor treatment (Fig. 6C), and the ranking of RAF inhibitors at the MOA-level (#1) was improved compared to the ranking of vemurafenib alone (#35) (Supp. Fig. 1B). Additionally, inhibitors of HDAC, EGFR, CHK, and SYK were identified as possibly beneficial for tumors with acquired resistance to RAF inhibition. Conversely, DMEA identified that drugs inhibiting MEK, MDM, nerve growth factor receptor, and HMGCR may be toxic towards tumors which are sensitive to RAF inhibitors (Fig. 6C). These results demonstrate that DMEA can amplify on-target signal to identify acquired resistance in tumors and other drug MOAs which may be beneficial based on patient biopsies.

### DMEA identifies potential senescence-inducing and senolytic drug MOAs for primary human mammary epithelial cells

Lastly, we sought to demonstrate how DMEA can be used as a discovery tool. As an example, we analyzed our recently published proteomic signature of replicative senescence, a stress-induced irreversible growth arrest associated with aging, in primary human mammary epithelial cells (HMECs) [39]. To highlight the versatility of DMEA to either identify similar or selectively toxic drug MOA, we analyzed the same molecular signature using either the CMap L1000 Query or our WGV-based approach to rank drugs, respectively (Fig. 7A). First, we performed a CMap L1000 Query using the gene names for the up- and down-regulated proteins to predict drug MOAs that could induce senescence in HMECs. Using the CMap results, DMEA revealed positive enrichment for MOAs including proteasome, HDAC, HMGCR, and MDM inhibitors (Fig. 7B), suggesting that treatment with drugs from these MOAs may induce senescence in primary HMECs. Among the MOAs with significant negative enrichments were Na/K-ATPase inhibitors and matrix metalloprotease inhibitors, suggesting that these drug MOAs might antagonize senescence in primary HMECs.

**Figure 7.**
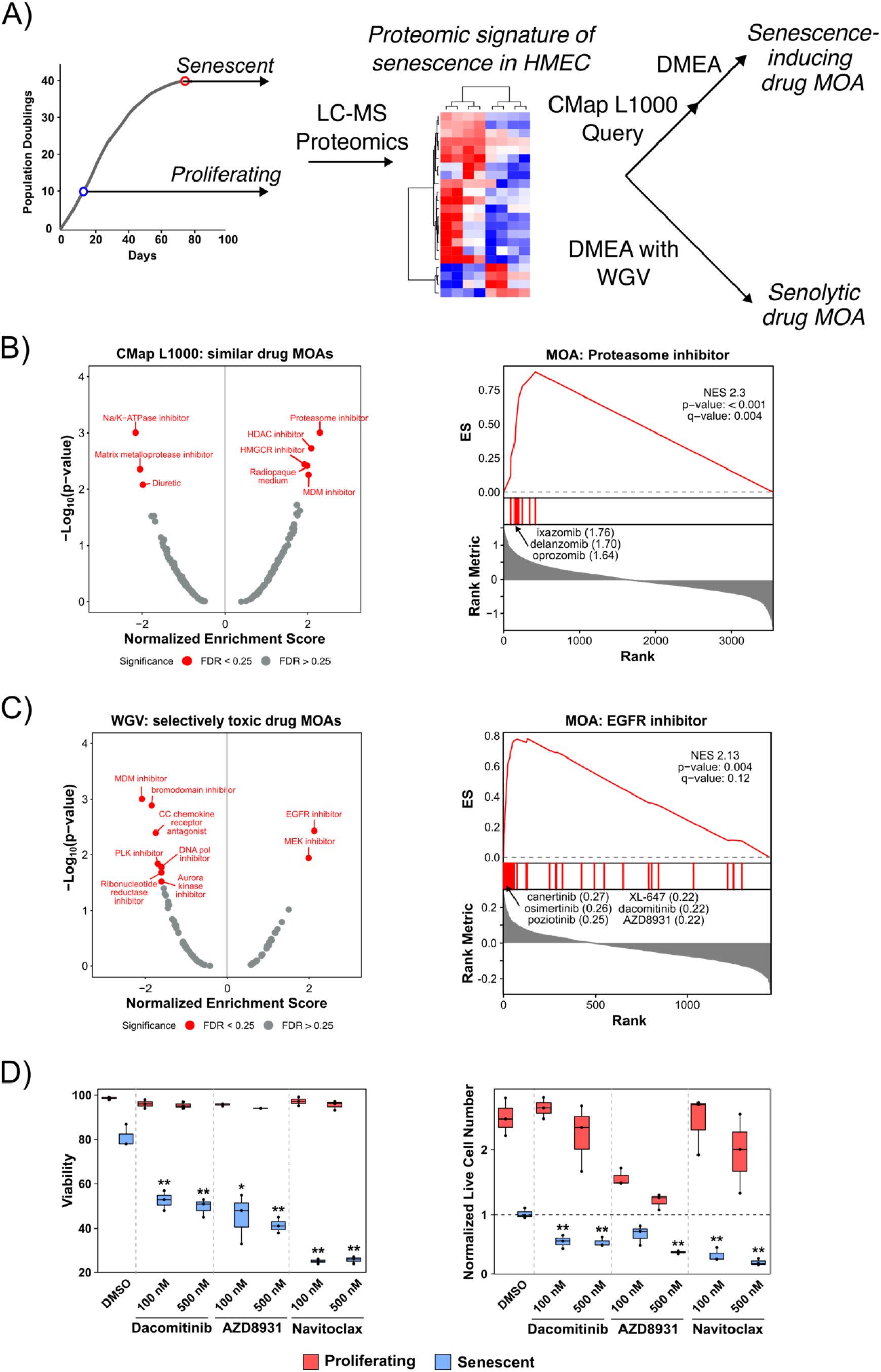
DMEA identifies potential senescence-inducing and senolytic drug MOAs for primary HMECs. A) Schematic detailing how the proteomic signature of replicative senescence in primary HMECs [39] was used to identify either senescence-inducing or senolytic drug MOAs. B) DMEA results for senescence-inducing drug MOAs. (Left) Volcano plot of NES versus −log_10_(p-value) for drug MOAs from DMEA. Red text indicates MOAs with p-value < 0.05 and FDR < 0.25. (Right) Mountain plot showing the positive enrichment of the proteasome inhibitor MOA in the rank-ordered drug list of CMap L1000 connectivity scores. The proteasome inhibitors with the most positive connectivity scores are highlighted. C) DMEA results for senolytic drug MOAs. (Left) Volcano plot of NES versus −log_10_(p-value) for drug MOAs from DMEA. Red text indicates MOAs with p-value < 0.05 and FDR < 0.25. (Right) Mountain plot showing the negative enrichment of the EGFR inhibitor MOA in the rank-ordered drug list of correlation coefficients. The EGFR inhibitors with the most negative correlation coefficients are highlighted. D) The EGFR inhibitors dacomitinib and AZD8931 and the senolytic compound navitoclax exhibited senolytic activity in HMECs. Proliferating HMECs (PD ∼12) were treated with DMSO or 2 μM triapine for 3 days to induce proliferating or senescent phenotypes, respectively, as in our previous work [45]. Proliferating and senescent HMECs were then treated with DMSO (negative control), 100 nM / 500 nM dacomitinib, 100 nM / 500 nM AZD8931, or 100 / 500 nM navitoclax for 3 days, after which cell viability and live cell number were measured by trypan blue staining. The live cell number was normalized to the number of live cells present at the time of drug treatment. * and ** represent p < 0.05 and 0.01, respectively, compared to the senescent DMSO control calculated by Student’s t-test.

Second, we analyzed the same proteomic signature of senescence with our own WGV-based method to identify selectively toxic MOAs based on the input molecular signature (Supp. Fig. 3). In contrast to analysis of the CMap L1000 Query which identified MOAs which induce similar gene expression, this DMEA pipeline instead predicted drug MOAs that may be toxic to cells with similar gene expression profiles as senescent HMEC (i.e., senolytic MOAs). Using this approach, we found that the EGFR and MEK inhibitor MOAs were significantly positively enriched in the rank-ordered drug list (Fig. 7C). This suggests that compounds from these MOAs may be senolytic in HMECs. Among the negatively enriched drug MOAs which may be more toxic to non-senescent, proliferating HMECs, we found MDM inhibitors, bromodomain inhibitors, and other MOAs.

Next, we experimentally tested the hypothesis that EGFR inhibitors exhibit selective toxicity against senescent HMECs using the same cells with which we generated the proteomic signature of senescence [45]. We treated proliferating and senescent HMECs with DMSO (negative control), the EGFR inhibitor dacomitinib, the EGFR inhibitor AZD8931, or the known senolytic compound navitoclax (positive control) at 100 and 500 nM. After 3 days of treatment, dacomitinib, AZD8931, and navitoclax significantly reduced the viability of senescent but not proliferating HMECs at both concentrations (Fig. 7D). Additionally, all three drugs reduced the total number of viable senescent cells. Notably, low concentrations of dacomitinib and navitoclax (100 nM) were selectively toxic to senescent cells without affecting the growth rate of non-senescent, proliferating HMECs. In contrast, although 100 nM of AZD8931 was selectively toxic to senescent HMEC, AZD8931 also reduced the growth rate of proliferating, non-senescent HMECs. These experimental results support that the EGFR inhibitors dacomitinib and AZD8931 are novel senolytic compounds in HMECs, validating a hypothesis generated by DMEA.

## DISCUSSION

Here, we introduce Drug Mechanism Enrichment Analysis (DMEA), a user-friendly bioinformatic method to better prioritize drug candidates for repurposing by grouping drugs based on shared MOA. Similar to how GSEA enhances biological interpretation of transcriptomic data [32], DMEA improves drug repurposing by aggregating information across many drugs with a common MOA instead of considering each drug independently. We have demonstrated the power and sensitivity of DMEA first with simulated data (Fig. 3; Supp. Fig. 4) and then with real examples including gene expression connectivity scores (Fig. 4), cell viability connectivity scores (Fig. 5), and weighted gene voting molecular classification scores (Fig. 6-7). In all cases, DMEA ranked the true drug MOA sensitivity or similarity higher than the original ranking of the single-drug agent (Supp. Fig. 1). In addition, DMEA improves upon existing tools for analyzing enriched MOA in drug lists in terms of flexibility, statistical rigor, and visual outputs (Fig. 1). This demonstrates that DMEA helps better prioritize drug treatments by improving the on-target identification of candidate drugs.

Importantly, our results demonstrate the ability of DMEA to analyze a variety of input rank-ordered drug lists from different drug repurposing algorithms to identify enrichment of diverse MOAs (e.g., kinase inhibitors, proteasome inhibitors, metabolic pathway inhibitors). In these validation cases, DMEA not only identified the expected drug MOAs (e.g., EGFR inhibitor MOA given a signature of EGFR inhibitor resistance), but also MOAs which may exhibit toxicity against tumor cells resistant to the input signature of interest. One interesting example is that DMEA identified HMGCR inhibitors as potentially toxic to cancer cells with intrinsic resistance to EGFR inhibitors (Fig. 6A-B; Supp. Fig. 5). Indeed, this finding is supported by published work demonstrating that HMGCR inhibitors can overcome resistance to EGFR inhibitors in NSCLC cells by inhibiting AKT [46, 47]. In addition, DMEA also identified that EGFR inhibitor-resistant cells may be sensitive to MDM inhibitors (Fig. 6A-B; Supp. Fig. 5), a finding that is supported by published work showing that MDM2 mediates resistance to EGFR inhibitors in mouse models of NSCLC [48]. Furthermore, our analysis suggested that melanomas sensitive to BRAF inhibitors may also be sensitive to MEK inhibitors (Fig. 6C), an observation that is supported by clinical trials showing that combination treatment with BRAF and MEK inhibitors is more effective than inhibition of BRAF alone in BRAF-mutant melanoma patients [49]. Finally, for melanomas with acquired resistance to RAF inhibitors, DMEA identified CHK inhibitors and SYK inhibitors as potentially beneficial. In fact, both CHK1 and SYK kinases have been identified as drug targets for melanomas resistant to RAF inhibitors [50, 51]. Collectively, these results support that DMEA can even identify drug mechanisms beneficial for combination treatments and drug-resistant cancers.

To demonstrate the power of DMEA for biological discovery, we analyzed our recently published proteomic signature of replicative senescence in primary HMECs [39] (Fig. 7). To illustrate the difference between CMap and DMEA’s interpretation of an input gene signature, we used: 1) the CMap L1000 Query followed by DMEA to identify similar (e.g., senescence-inducing) drug MOA and 2) DMEA with WGV molecular classification scores to identify selectively toxic (e.g., senolytic) drug MOA. Both senescence-inducing and senolytic compounds have great therapeutic promise in aging [52–56], cancer [57, 58], and other diseases [59]. Among the potential senescence-inducing drug MOAs we identified were proteasome, HDAC, HMGCR, and MDM inhibitors (Fig. 7B). Indeed, experimental evidence has shown that proteasome inhibitors induce senescence in primary fibroblasts [60, 61] and that HDAC inhibitors can induce senescence in cancer cells [62–64]. For potential senolytic MOAs, we identified EGFR and MEK inhibitors (Fig. 7C), and we subsequently experimentally validated the senolytic activity of the EGFR inhibitors dacomitinib and AZD8931 in a drug-induced model of HMEC senescence (Fig. 7D). To our knowledge, this is the first demonstration that EGFR inhibitors can exhibit senolytic activity. Although we did not test whether MEK inhibitors would exhibit senolytic activity in primary HMECs, it has been shown in Ras-expressing cells that MEK inhibitors selectively kill senescent cells [65]. Taken together, our results indicate that DMEA is a powerful tool for repurposing drug MOAs based on selectivity against or similarity to a given molecularly characterized cell state.

Despite the success of our validation examples, we note that DMEA is limited by our knowledge of drug MOA. For many targeted therapeutics, the putative MOA may be incorrect [31]. Nevertheless, DMEA mitigates the risk of false positives by evaluating groups of drugs which share a MOA rather than relying on results from individual drugs alone. Thus, even if some drugs are misannotated, DMEA may still correctly identify enriched MOAs by aggregating information across multiple drugs, rather than considering each drug independently. However, improved annotation of drug MOAs and targets, potentially through newer approaches including metabolomics [66] and proteomics [67], will improve the power of DMEA.

In summary, Drug Mechanism Enrichment Analysis (DMEA) improves prioritization of drugs for repurposing by grouping drugs that share mechanisms of action (MOAs). DMEA can thus be used to further process rank-ordered lists of drugs from drug repurposing algorithms to sharpen on-target signal. To provide an easily accessible tool for drug repurposing, we also added the option to pair DMEA with WGV molecular classification as well as public databases of transcriptomic profiles (e.g., L1000, CCLE) and drug screens (e.g., PRISM). With this feature, DMEA can interpret an input gene signature to rapidly identify drug mechanisms which exhibit selective toxicity towards cell states (e.g., cancer, senescence). Furthermore, our results support that DMEA has potential to aid in the discovery of therapeutics for combination treatments or drug-resistant cancers. DMEA is publicly available to use either as a web application or an R package at https://belindabgarana.github.io/DMEA.

## DECLARATIONS

### ETHICS APPROVAL AND CONSENT TO PARTICIPATE

Not applicable.

### CONSENT FOR PUBLICATION

Not applicable.

### AVAILABILITY OF DATA AND MATERIALS

All data and code are publicly available at https://github.com/BelindaBGarana/DMEA.

### AUTHORS’ CONTRIBUTIONS

BBG, JHJ, and NAG conceived the project. BBG and JHJ wrote the R code to implement the project. BBG performed the computational experiments and downstream bioinformatics and developed the Shiny web application and R package. AD designed, conducted, and performed data analysis for the wet lab experiments with senescent cells. HH designed and developed the website hosted on GitHub. NAG supervised the project and provided funding. BBG and NAG wrote the manuscript. All authors read and approved the final manuscript.

## ACKNOWLEDGMENTS

Not applicable.

## REFERENCES

1. Li H, Ning S, Ghandi M, Kryukov GV, Gopal S, Deik A, et al. The landscape of cancer cell line metabolism. Nature Medicine. 2019;25:850.

2. Nusinow DP, Szpyt J, Ghandi M, Rose CM, McDonald ER, Kalocsay M, et al. Quantitative Proteomics of the Cancer Cell Line Encyclopedia. Cell. 2020;180:387–402.e16.

3. Ghandi M, Huang FW, Jané-Valbuena J, Kryukov GV, Lo CC, McDonald ER, et al. Next-generation characterization of the Cancer Cell Line Encyclopedia. Nature. 2019;569:503–8.

4. Frejno M, Meng C, Ruprecht B, Oellerich T, Scheich S, Kleigrewe K, et al. Proteome activity landscapes of tumor cell lines determine drug responses. Nature Communications. 2020;11:3639.

5. Corsello SM, Nagari RT, Spangler RD, Rossen J, Kocak M, Bryan JG, et al. Discovering the anticancer potential of non-oncology drugs by systematic viability profiling. Nat Cancer. 2020;1:235–48.

6. Behan FM, Iorio F, Picco G, Gonçalves E, Beaver CM, Migliardi G, et al. Prioritization of cancer therapeutic targets using CRISPR–Cas9 screens. Nature. 2019;568:511–6.

7. Meyers RM, Bryan JG, McFarland JM, Weir BA, Sizemore AE, Xu H, et al. Computational correction of copy number effect improves specificity of CRISPR–Cas9 essentiality screens in cancer cells. Nat Genet. 2017;49:1779–84.

8. Subramanian A, Narayan R, Corsello SM, Peck DD, Natoli TE, Lu X, et al. A Next Generation Connectivity Map: L1000 Platform and the First 1,000,000 Profiles. Cell. 2017;171:1437–1452.e17.

9. Parca L, Pepe G, Pietrosanto M, Galvan G, Galli L, Palmeri A, et al. Modeling cancer drug response through drug-specific informative genes. Sci Rep. 2019;9:15222.

10. Wei D, Liu C, Zheng X, Li Y. Comprehensive anticancer drug response prediction based on a simple cell line-drug complex network model. BMC Bioinformatics. 2019;20:44.

11. Suphavilai C, Bertrand D, Nagarajan N. Predicting Cancer Drug Response using a Recommender System. Bioinformatics. 2018;34:3907–14.

12. Yuan H, Paskov I, Paskov H, González AJ, Leslie CS. Multitask learning improves prediction of cancer drug sensitivity. Sci Rep. 2016;6:31619.

13. Choi J, Park S, Ahn J. RefDNN: a reference drug based neural network for more accurate prediction of anticancer drug resistance. Sci Rep. 2020;10:1861.

14. Gonçalves E, Segura-Cabrera A, Pacini C, Picco G, Behan FM, Jaaks P, et al. Drug mechanism-of-action discovery through the integration of pharmacological and CRISPR screens. Molecular Systems Biology. 2020;16:e9405.

15. Nguyen L, Dang CC, Ballester PJ. Systematic assessment of multi-gene predictors of pan-cancer cell line sensitivity to drugs exploiting gene expression data. F1000Res. 2017;5:ISCB Comm J–2927.

16. Gao S, Han L, Luo D, Liu G, Xiao Z, Shan G, et al. Modeling drug mechanism of action with large scale gene-expression profiles using GPAR, an artificial intelligence platform. BMC Bioinformatics. 2021;22:17.

17. Napolitano F, Carrella D, Mandriani B, Pisonero-Vaquero S, Sirci F, Medina DL, et al. gene2drug: a computational tool for pathway-based rational drug repositioning. Bioinformatics. 2018;34:1498–505.

18. Fang M, Richardson B, Cameron CM, Dazard J-E, Cameron MJ. Drug perturbation gene set enrichment analysis (dpGSEA): a new transcriptomic drug screening approach. BMC Bioinformatics. 2021;22:22.

19. Prinz J, Vogt I, Adornetto G, Campillos M. A Novel Drug-Mouse Phenotypic Similarity Method Detects Molecular Determinants of Drug Effects. PLOS Computational Biology. 2016;12:e1005111.

20. Malyutina A, Majumder MM, Wang W, Pessia A, Heckman CA, Tang J. Drug combination sensitivity scoring facilitates the discovery of synergistic and efficacious drug combinations in cancer. PLOS Computational Biology. 2019;15:e1006752.

21. Gayvert KM, Aly O, Platt J, Bosenberg MW, Stern DF, Elemento O. A Computational Approach for Identifying Synergistic Drug Combinations. PLOS Computational Biology. 2017;13:e1005308.

22. Rivas-Barragan D, Mubeen S, Bernat FG, Hofmann-Apitius M, Domingo-Fernández D. Drug2ways: Reasoning over causal paths in biological networks for drug discovery. PLOS Computational Biology. 2020;16:e1008464.

23. Domingo-Fernández D, Gadiya Y, Patel A, Mubeen S, Rivas-Barragan D, Diana CW, et al. Causal reasoning over knowledge graphs leveraging drug-perturbed and disease-specific transcriptomic signatures for drug discovery. PLOS Computational Biology. 2022;18:e1009909.

24. Szalai B, Subramanian V, Holland CH, Alföldi R, Puskás LG, Saez-Rodriguez J. Signatures of cell death and proliferation in perturbation transcriptomics data—from confounding factor to effective prediction. Nucleic Acids Research. 2019;47:10010–26.

25. Napolitano F, Sirci F, Carrella D, di Bernardo D. Drug-set enrichment analysis: a novel tool to investigate drug mode of action. Bioinformatics. 2016;32:235–41.

26. Brum AM, van de Peppel J, van der Leije CS, Schreuders-Koedam M, Eijken M, van der Eerden BCJ, et al. Connectivity Map-based discovery of parbendazole reveals targetable human osteogenic pathway. Proceedings of the National Academy of Sciences. 2015;112:12711–6.

27. Zhang M, Luo H, Xi Z, Rogaeva E. Drug Repositioning for Diabetes Based on “Omics” Data Mining. PLOS ONE. 2015;10:e0126082.

28. Tsoi J, Robert L, Paraiso K, Galvan C, Sheu KM, Lay J, et al. Multi-stage Differentiation Defines Melanoma Subtypes with Differential Vulnerability to Drug-Induced Iron-Dependent Oxidative Stress. Cancer Cell. 2018;33:890–904.e5.

29. Viswanathan VS, Ryan MJ, Dhruv HD, Gill S, Eichhoff OM, Seashore-Ludlow B, et al. Dependency of a therapy-resistant state of cancer cells on a lipid peroxidase pathway. Nature. 2017;547:453–7.

30. Haslam A, Kim MS, Prasad V. Updated estimates of eligibility for and response to genome-targeted oncology drugs among US cancer patients, 2006-2020. Ann Oncol. 2021;32:926–32.

31. Lin A, Giuliano CJ, Palladino A, John KM, Abramowicz C, Yuan ML, et al. Off-target toxicity is a common mechanism of action of cancer drugs undergoing clinical trials. Science Translational Medicine. 2019;11.

32. Subramanian A, Tamayo P, Mootha VK, Mukherjee S, Ebert BL, Gillette MA, et al. Gene set enrichment analysis: A knowledge-based approach for interpreting genome-wide expression profiles. Proceedings of the National Academy of Sciences. 2005;102:15545–50.

33. Golub TR, Slonim DK, Tamayo P, Huard C, Gaasenbeek M, Mesirov JP, et al. Molecular Classification of Cancer: Class Discovery and Class Prediction by Gene Expression Monitoring. Science. 1999;286:531–7.

34. Maejima T, Inoue T, Kanki Y, Kohro T, Li G, Ohta Y, et al. Direct evidence for pitavastatin induced chromatin structure change in the KLF4 gene in endothelial cells. PLoS One. 2014;9:e96005.

35. Greger JG, Eastman SD, Zhang V, Bleam MR, Hughes AM, Smitheman KN, et al. Combinations of BRAF, MEK, and PI3K/mTOR inhibitors overcome acquired resistance to the BRAF inhibitor GSK2118436 dabrafenib, mediated by NRAS or MEK mutations. Mol Cancer Ther. 2012;11:909–20.

36. Wang Q, Mora-Jensen H, Weniger MA, Perez-Galan P, Wolford C, Hai T, et al. ERAD inhibitors integrate ER stress with an epigenetic mechanism to activate BH3-only protein NOXA in cancer cells. Proc Natl Acad Sci U S A. 2009;106:2200–5.

37. Narla A, Dutt S, McAuley JR, Al-Shahrour F, Hurst S, McConkey M, et al. Dexamethasone and lenalidomide have distinct functional effects on erythropoiesis. Blood. 2011;118:2296–304.

38. Brachmann SM, Kleylein-Sohn J, Gaulis S, Kauffmann A, Blommers MJJ, Kazic-Legueux M, et al. Characterization of the mechanism of action of the pan class I PI3K inhibitor NVP-BKM120 across a broad range of concentrations. Mol Cancer Ther. 2012;11:1747–57.

39. Delfarah A, Hartel NG, Zheng D, Yang J, Graham NA. Identification of a Proteomic Signature of Senescence in Primary Human Mammary Epithelial Cells. J Proteome Res. 2021;20:5169–79.

40. Ritchie ME, Phipson B, Wu D, Hu Y, Law CW, Shi W, et al. limma powers differential expression analyses for RNA-sequencing and microarray studies. Nucleic Acids Research. 2015;43:e47–e47.

41. Hoeflich KP, O’Brien C, Boyd Z, Cavet G, Guerrero S, Jung K, et al. In vivo antitumor activity of MEK and phosphatidylinositol 3-kinase inhibitors in basal-like breast cancer models. Clin Cancer Res. 2009;15:4649–64.

42. Balko JM, Potti A, Saunders C, Stromberg A, Haura EB, Black EP. Gene expression patterns that predict sensitivity to epidermal growth factor receptor tyrosine kinase inhibitors in lung cancer cell lines and human lung tumors. BMC Genomics. 2006;7:289.

43. Coldren CD, Helfrich BA, Witta SE, Sugita M, Lapadat R, Zeng C, et al. Baseline Gene Expression Predicts Sensitivity to Gefitinib in Non–Small Cell Lung Cancer Cell Lines. Mol Cancer Res. 2006;4:521–8.

44. Kemper K, Krijgsman O, Kong X, Cornelissen-Steijger P, Shahrabi A, Weeber F, et al. BRAF(V600E) Kinase Domain Duplication Identified in Therapy-Refractory Melanoma Patient-Derived Xenografts. Cell Rep. 2016;16:263–77.

45. Delfarah A, Parrish S, Junge JA, Yang J, Seo F, Li S, et al. Inhibition of nucleotide synthesis promotes replicative senescence of human mammary epithelial cells. J Biol Chem. 2019;:jbc.RA118.005806.

46. Hwang K-E, Kwon S-J, Kim Y-S, Park D-S, Kim B-R, Yoon K-H, et al. Effect of simvastatin on the resistance to EGFR tyrosine kinase inhibitors in a non-small cell lung cancer with the T790M mutation of EGFR. Experimental Cell Research. 2014;323:288–96.

47. Chen J, Bi H, Hou J, Zhang X, Zhang C, Yue L, et al. Atorvastatin overcomes gefitinib resistance in KRAS mutant human non-small cell lung carcinoma cells. Cell Death Dis. 2013;4:e814–e814.

48. Shen H, Wang G-C, Li X, Ge X, Wang M, Shi Z-M, et al. S6K1 blockade overcomes acquired resistance to EGFR-TKIs in non-small cell lung cancer. Oncogene. 2020;39:7181–95.

49. Long GV, Stroyakovskiy D, Gogas H, Levchenko E, de Braud F, Larkin J, et al. Combined BRAF and MEK inhibition versus BRAF inhibition alone in melanoma. N Engl J Med. 2014;371:1877–88.

50. Hwang B-J, Adhikary G, Eckert RL, Lu A-L. Chk1 inhibition as a novel therapeutic strategy in melanoma. Oncotarget. 2018;9:30450–64.

51. Abecunas C, Whitehead CE, Ziemke EK, Baumann DG, Frankowski-McGregor CL, Sebolt-Leopold JS, et al. Loss of NF1 in Melanoma Confers Sensitivity to SYK Kinase Inhibition. Cancer Research. 2023;:OF1–16.

52. Xu M, Pirtskhalava T, Farr JN, Weigand BM, Palmer AK, Weivoda MM, et al. Senolytics improve physical function and increase lifespan in old age. Nature Medicine. 2018;24:1246–56.

53. Baar MP, Brandt RMC, Putavet DA, Klein JDD, Derks KWJ, Bourgeois BRM, et al. Targeted Apoptosis of Senescent Cells Restores Tissue Homeostasis in Response to Chemotoxicity and Aging. Cell. 2017;169:132–147.e16.

54. Baker DJ, Wijshake T, Tchkonia T, LeBrasseur NK, Childs BG, van de Sluis B, et al. Clearance of p16Ink4a-positive senescent cells delays ageing-associated disorders. Nature. 2011;479:232–6.

55. Baker DJ, Childs BG, Durik M, Wijers ME, Sieben CJ, Zhong J, et al. Naturally occurring p16 Ink4a-positive cells shorten healthy lifespan. Nature. 2016;530:184–9.

56. Cai Y, Zhou H, Zhu Y, Sun Q, Ji Y, Xue A, et al. Elimination of senescent cells by β-galactosidase-targeted prodrug attenuates inflammation and restores physical function in aged mice. Cell Research. 2020;30:574–89.

57. Dörr JR, Yu Y, Milanovic M, Beuster G, Zasada C, Däbritz JHM, et al. Synthetic lethal metabolic targeting of cellular senescence in cancer therapy. Nature. 2013;501:421–5.

58. Guerrero A, Herranz N, Sun B, Wagner V, Gallage S, Guiho R, et al. Cardiac glycosides are broad-spectrum senolytics. Nat Metab. 2019;1:1074–88.

59. Hickson LJ, Langhi Prata LGP, Bobart SA, Evans TK, Giorgadze N, Hashmi SK, et al. Senolytics decrease senescent cells in humans: Preliminary report from a clinical trial of Dasatinib plus Quercetin in individuals with diabetic kidney disease. EBioMedicine. 2019;47:446–56.

60. Chondrogianni N, Gonos ES. Proteasome inhibition induces a senescence-like phenotype in primary human fibroblasts cultures. Biogerontology. 2004;5:55–61.

61. Torres C, Lewis L, Cristofalo VJ. Proteasome inhibitors shorten replicative life span and induce a senescent-like phenotype of human fibroblasts. Journal of Cellular Physiology. 2006;207:845–53.

62. Lorenz V, Hessenkemper W, Rödiger J, Kyrylenko S, Kraft F, Baniahmad A. Sodium butyrate induces cellular senescence in neuroblastoma and prostate cancer cells. Horm Mol Biol Clin Investig. 2011;7:265–72.

63. Venkatesh R, Ramaiah MJ, Gaikwad HK, Janardhan S, Bantu R, Nagarapu L, et al. Luotonin-A based quinazolinones cause apoptosis and senescence via HDAC inhibition and activation of tumor suppressor proteins in HeLa cells. Eur J Med Chem. 2015;94:87–101.

64. Vargas JE, Filippi-Chiela EC, Suhre T, Kipper FC, Bonatto D, Lenz G. Inhibition of HDAC increases the senescence induced by natural polyphenols in glioma cells. Biochem Cell Biol. 2014;92:297–304.

65. Kochetkova EY, Blinova GI, Bystrova OA, Martynova MG, Pospelov VA, Pospelova TV. Targeted elimination of senescent Ras-transformed cells by suppression of MEK/ERK pathway. Aging (Albany NY). 2017;9:2352–75.

66. Holbrook-Smith D, Durot S, Sauer U. High-throughput metabolomics predicts drug-target relationships for eukaryotic proteins. Mol Syst Biol. 2022;18:e10767.

67. Savitski MM, Reinhard FBM, Franken H, Werner T, Savitski MF, Eberhard D, et al. Tracking cancer drugs in living cells by thermal profiling of the proteome. Science. 2014. https://doi.org/10.1126/science.1255784.

