## Supplemental Figures for "Drug mechanism enrichment analysis improves prioritization of therapeutics for repurposing"

### DATA AVAILABILITY

All code and data used in this manuscript, as well as a web application and an R package to run DMEA, are available at https://belindabgarana.github.io/DMEA.

### SUPPLEMENTARY FIGURES

A)

| *Using CMap L1000 Query Tool* | | Rank / List Length | | | |
|---|---|---|---|---|---|
| | | **CMap L1000 Query** | | | **DMEA** |
| **Dataset** | **Drug (MOA)** | Drug | MOA | PCL | MOA |
| GSE32547 | Pitavastatin (HMGCR inhibitor) | 24 / 3868 | 1 / 234 | 1 / 57 | 1 / 177 |
| GSE35230 | GSK212 (MEK inhibitor) | NF / 3868 | 1 / 234 | 1 / 57 | 1 / 179 |
| GSE14003 | Bortezomib (Proteasome inhibitor) | 14 / 3868 | 1 / 234 | 3 / 57 | 1 / 180 |
| GSE28896 | Dexamethasone (Glucocorticoid agonist) | 390 / 3868 | NF / 234 | 2 / 57 | 2 / 177 |
| GSE33643 | BEZ235 (mTOR & PI3K inhibitor) | 24 / 3868 | 1 & NF / 234 | 1 & 2 / 57 | 1 & 2 / 183 |

B)

| *Using Other Drug Repurposing Algorithms* | | Rank / List Length | | |
|---|---|---|---|---|
| | | **Input** | | **DMEA** |
| **Dataset** | **Drug (MOA)** | Algorithm | Drug | MOA |
| PRISM | Lovastatin (HMGCR inhibitor) | CMap PRISM Query | 3 / 1174 | 1 / 65 |
| GSE31625 | Erlotinib (EGFR inhibitor) | WGV correlation with PRISM drug AUC | 16 / 1351 | 1 / 85 |
| GSE12790 | | | 10 / 1351 | 1 / 85 |
| Coldren et al. | Gefitinib (EGFR inhibitor) | | 13 / 1351 | 1 / 85 |
| GSE66539 | Vemurafenib (RAF inhibitor) | | 35 / 1351 | 1 / 85 |

**Supp. Figure 1. DMEA generates MOA rankings that are improved over single-drug rankings and is compatible with more datasets than the CMap Query tool.** A) Comparison of DMEA's MOA rankings to rankings generated by the CMap L1000 Query. DMEA's rankings are the same or improved compared to CMap's MOA or PCL rankings in all cases and improved compared to the single-drug rankings in all cases. NF: not found because this drug is not present in the L1000 database. B) DMEA's MOA rankings also improve upon single-drug rankings in other cases where CMap cannot provide MOA or PCL analysis.

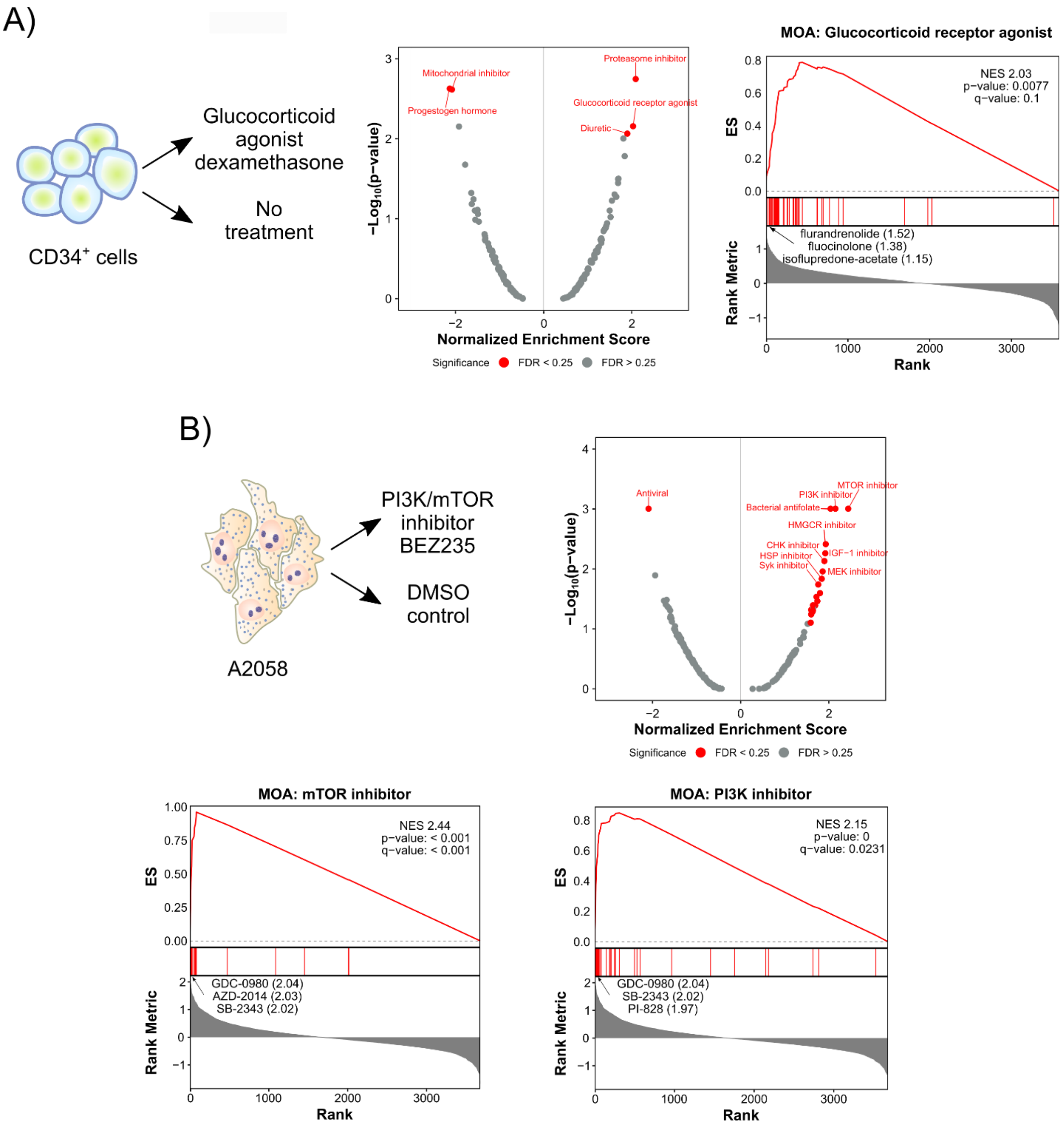


**Supp. Figure 2. DMEA identifies similar MOAs based on gene expression connectivity scores.** Rank-ordered drug lists were generated by querying the CMap L1000 gene expression perturbation signatures and then analyzed by DMEA. A) Human CD34+ cells treated with the glucocorticoid agonist dexamethasone (24 h) or untreated [1]; B) A2058 cells treated with the PI3K/MTOR inhibitor BEZ235 or DMSO [2]. Volcano plots summarizing the NES and -$\log_{10}$(p-value) for all tested drug MOAs and mountain plots of the expected MOAs are shown. Red text indicates MOAs with p-value < 0.05 and FDR < 0.25.

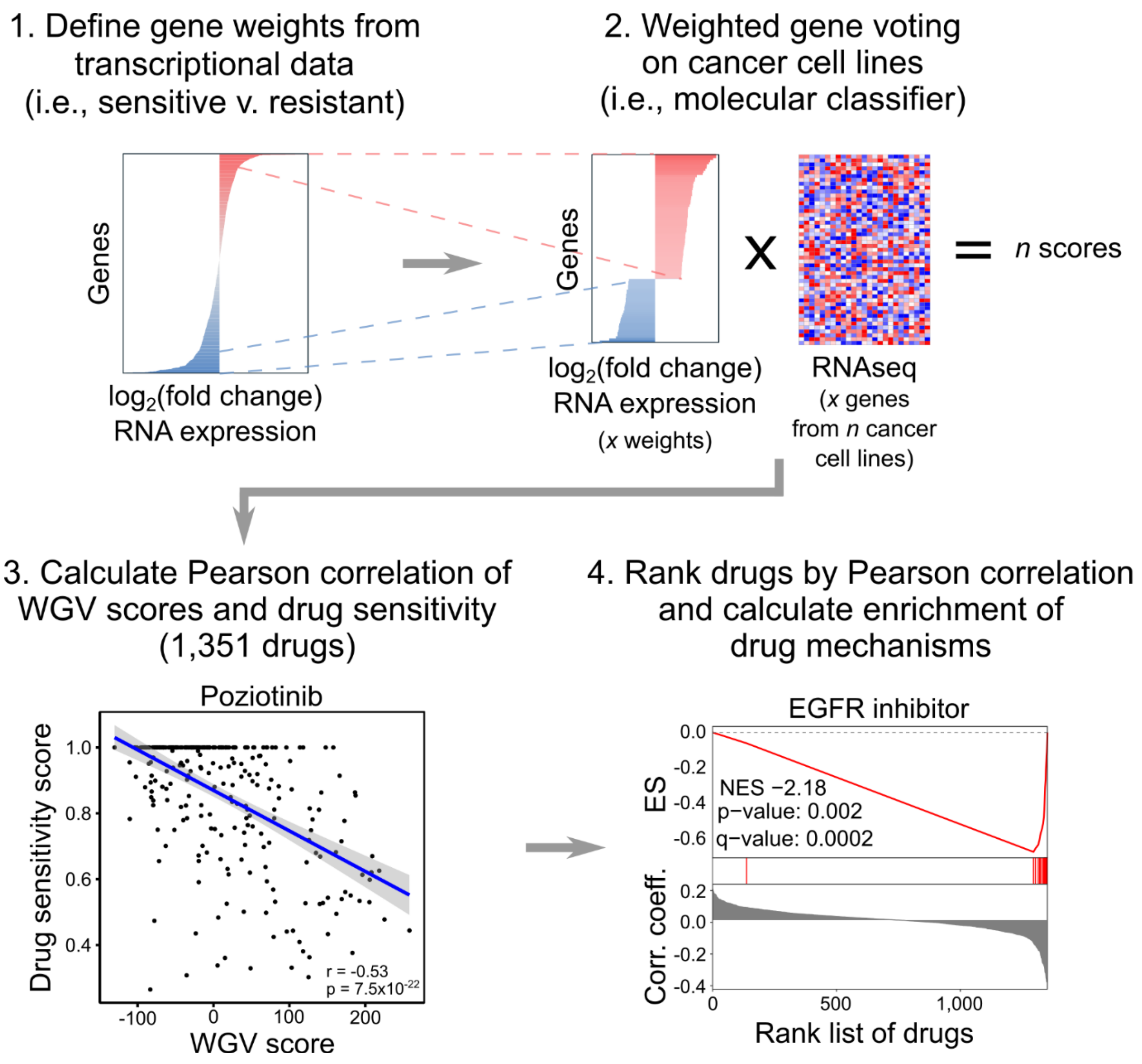


**Supp. Figure 3. Overview of Drug Mechanism Enrichment Analysis using WGV molecular classification scores.** DMEA can accept an input gene signature when paired with a molecular classification method such as weighted gene voting (WGV), correlations, and large public databases of gene expression and drug screens. First, gene weights are calculated as $\log_2$(fold change) values calculated between two groups of samples from transcriptional data. Only the *x* genes with $\log_2$(fold change) with q-value < 0.05 are selected for subsequent steps (maximum 500 genes). Second, the WGV molecular classifier score is calculated as the dot product between *x* gene weights and gene expression values from cancer cell lines (*x* genes by *n* cell lines). Here, we used 327 adherent cancer cell lines from the CCLE. Third, the Pearson correlation is calculated between the WGV score and drug sensitivity score (i.e., area under the curve (AUC) of cell viability versus drug concentration from the PRISM database) for each of the 1,351 drugs. An example is shown for the EGFR inhibitor poziotinib. Fourth, the 1,351 drugs are ranked by Pearson correlation coefficient, and the ranked list is analyzed using a GSEA-like algorithm to identify drug mechanisms that are enriched at either end of the ranked list. In total, 85 mechanisms-of-action are tested (Supp. Table 1). An example for EGFR inhibitors is shown. Each red tick mark represents a drug annotated as an EGFR inhibitor in the waterfall plot of correlation coefficients.

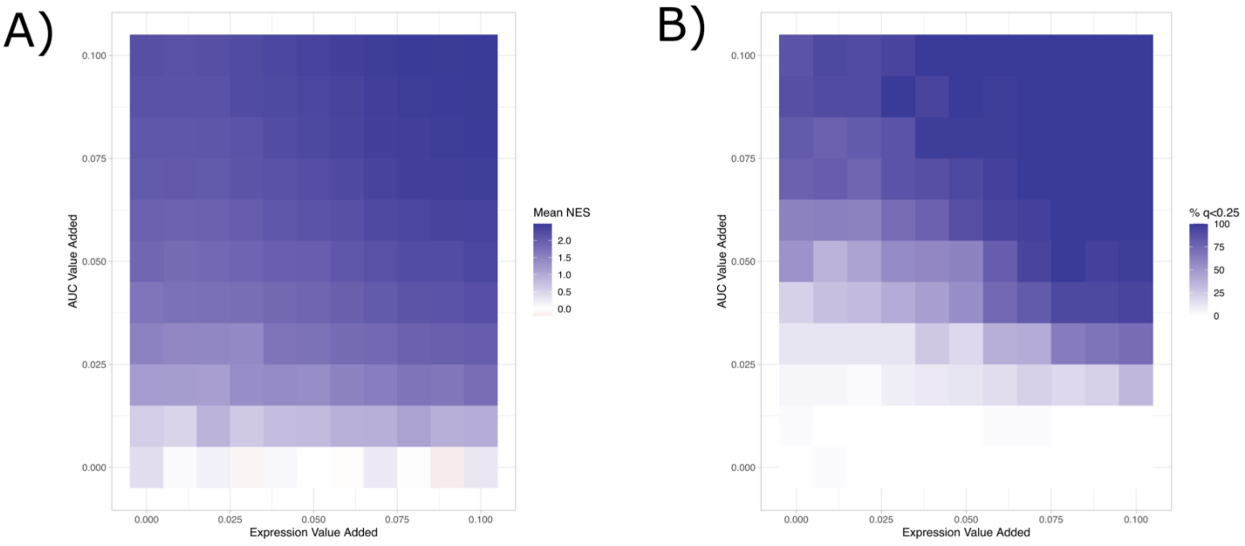


**Supp. Figure 4. Sensitivity analysis of DMEA pipeline with WGV using synthetic data.** Synthetic gene expression and drug sensitivity data was simulated with varying perturbations to RNA expression (x-axis) and drug sensitivity score (i.e., AUC, y-axis) (see Methods). A) Heatmap showing the average DMEA NES from 50 simulations for varying perturbations of the drug AUC and gene expression data. B) Heatmap showing the percent of DMEA replicates with FDR q-value < 0.05 from 50 simulations for varying perturbations of the drug AUC and gene expression data.

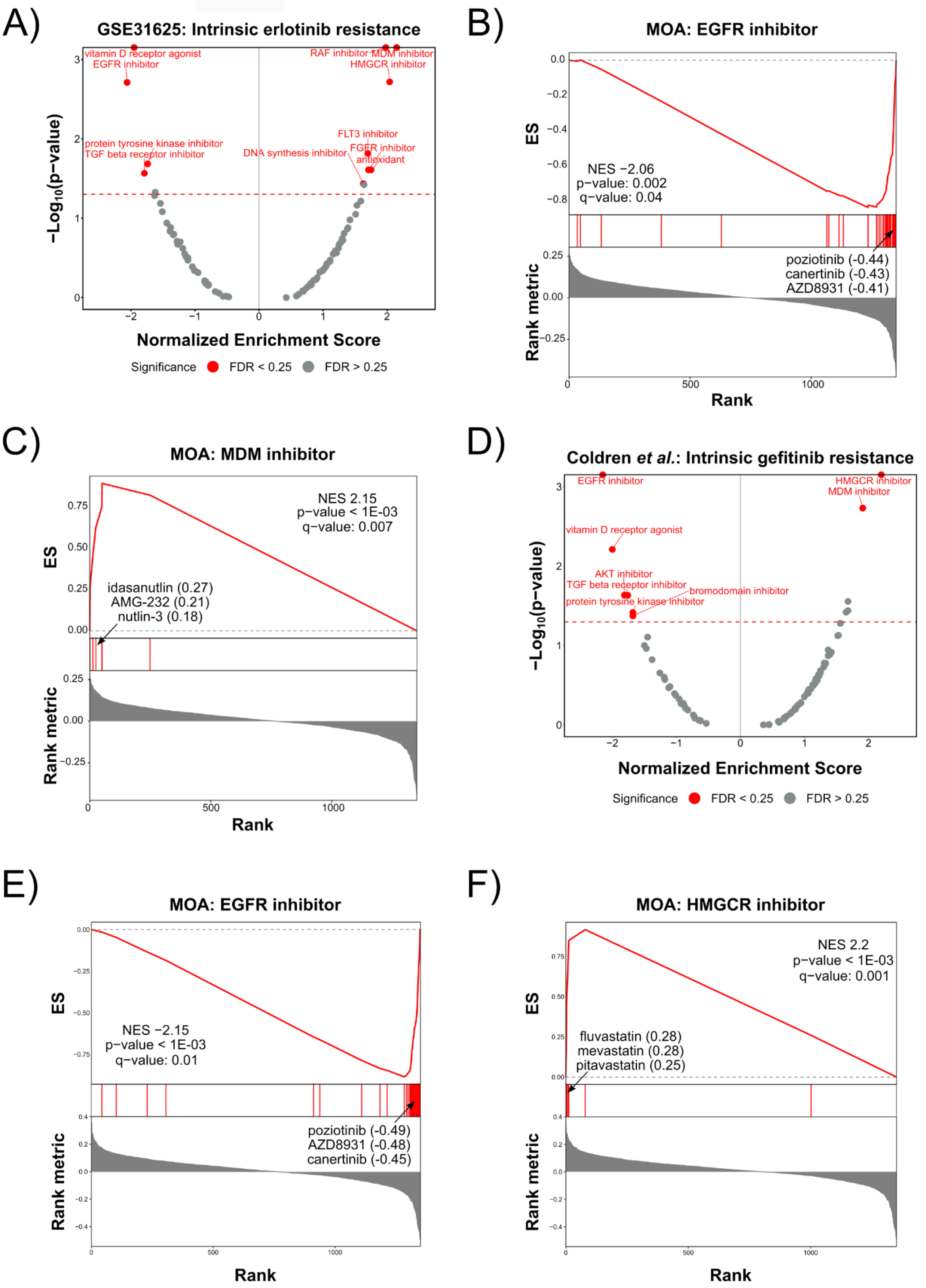
A)
GSE31625: Intrinsic erlotinib resistance
vitamin D receptor agonist
EGFR inhibitor
RAF inhibitor
MDM inhibitor
HMGCR inhibitor
protein tyrosine kinase inhibitor
TGF beta receptor inhibitor
FLT3 inhibitor
FGFR inhibitor
DNA synthesis inhibitor
antioxidant
−Log10(p−value)
Normalized Enrichment Score
Significance
FDR < 0.25
FDR > 0.25
B)
MOA: EGFR inhibitor
ES
NES −2.06
p−value: 0.002
q−value: 0.04
poziotinib (-0.44)
canertinib (-0.43)
AZD8931 (-0.41)
Rank metric
Rank
C)
MOA: MDM inhibitor
NES 2.15
p−value < 1E-03
q−value: 0.007
idasanutlin (0.27)
AMG-232 (0.21)
nutlin-3 (0.18)
ES
Rank metric
Rank
D)
Coldren et al.: Intrinsic gefitinib resistance
EGFR inhibitor
HMGCR inhibitor
MDM inhibitor
vitamin D receptor agonist
AKT inhibitor
TGF beta receptor inhibitor
bromodomain inhibitor
protein tyrosine kinase inhibitor
−Log10(p−value)
Normalized Enrichment Score
Significance
FDR < 0.25
FDR > 0.25
E)
MOA: EGFR inhibitor
ES
NES −2.15
p−value < 1E-03
q−value: 0.01
poziotinib (-0.49)
AZD8931 (-0.48)
canertinib (-0.45)
Rank metric
Rank
F)
MOA: HMGCR inhibitor
NES 2.2
p−value < 1E-03
q−value: 0.001
fluvastatin (0.28)
mevastatin (0.28)
pitavastatin (0.25)
ES
Rank metric
Rank

**Supp. Figure 5. DMEA identifies enrichment of the EGFR inhibitor MOA and other MOAs using external signatures of intrinsic EGFR inhibitor resistance.** Using gene expression signatures of intrinsic erlotinib (GSE31625 [3]) and gefitinib resistance (Coldren *et al.* [4]), we calculated WGV scores for 327 adherent cancer cell lines in the CCLE database. For each signature, the WGV scores were correlated with drug sensitivity data (i.e., AUC) for 1,351 drugs from the PRISM database. Drugs were then ranked by correlation coefficient, and DMEA was performed to identify enriched MOAs at either end of the rank-ordered drug list. A) Volcano plot of NES versus -$\log_{10}$(p-value) for DMEA using the GSE31625 signature of erlotinib resistance. Red text indicates MOAs with p-value < 0.05 and FDR < 0.25. B) Mountain plot showing that DMEA identified the EGFR inhibitor MOA as negatively enriched in the rank-ordered drug list generated using the GSE31625 signature of erlotinib resistance. The most negatively correlated EGFR inhibitors are highlighted along with their correlation coefficients. C) Mountain plot showing that DMEA identified the MDM inhibitor MOA as positively enriched in the rank-ordered drug list generated using the GSE31625 signature of erlotinib resistance. The most positively correlated MDM inhibitors are highlighted along with their correlation coefficients. D) Volcano plot of NES versus -$\log_{10}$(p-value) for DMEA using the Coldren *et al.* signature of gefitinib resistance. Red text indicates MOAs with p-value < 0.05 and FDR < 0.25. E) Mountain plot showing that DMEA identified the EGFR inhibitor MOA as negatively enriched in the rank-ordered drug list generated using the Coldren *et al.* signature of gefitinib resistance. The most negatively correlated EGFR inhibitors are highlighted along with their correlation coefficients. F) Mountain plot showing that DMEA identified the HMGCR inhibitor MOA as positively enriched in the rank-ordered drug list generated using the Coldren *et al.* signature of gefitinib resistance. The most positively correlated HMGCR inhibitors are highlighted along with their correlation coefficients.
